# Spatially resolved contrastive analysis of perturbation-driven microenvironmental heterogeneity across tissues and conditions

**DOI:** 10.64898/2025.12.08.693051

**Authors:** Yan Cui, Jasmine Blandin, Kipp Weiskopf, Na Sun

## Abstract

Understanding how tissues remodel in response to perturbations requires computational tools that can untangle condition-specific changes from the conserved tissue architecture. We present Haruka, a spatially aware contrastive learning framework that identifies salient (condition-specific) and background (shared) spatial domains across tissue slices and experimental conditions. Haruka integrates contrastive variational inference with an auxiliary microenvironment reconstruction task, enabling the model to learn spatial-context-informed embeddings that capture both perturbation effects and local neighborhood context. Benchmarking on simulated and real datasets demonstrates superior detection of spatially heterogeneous perturbation responses compared to existing methods. Applied to diverse spatial omics platforms, Haruka distinguished immunotherapy responders in melanoma, traced fibrosis progression in human lung tissue, and mapped treatment-resistant microenvironments in KRAS^G12D^-mutated lung cancer. Extended to spatially resolved multiomics data co-profiling transcriptomics and chromatin accessibility from an LPC-induced neuroinflammation mouse model, using a modality-parallel design that independently models each omics layer before integration, Haruka resolved condition-specific regulatory programs encoded across chromatin and transcriptome layers, linking spatial domain identity to transcription factor network architecture and human psychiatric disease genetics. Thus, Haruka provides a generalizable framework for spatial contrastive analysis, enabling systematic dissection of tissue organization, cellular plasticity, and microenvironmental remodeling across disease contexts and therapeutic perturbations.

## Introduction

Advances in high-throughput spatial omics technologies have revolutionized our ability to study biological systems by capturing molecular information directly within intact tissue. By profiling multiple tissue slices in parallel, these methods enable detailed assessment of spatial heterogeneity and reproducibility across biological and experimental conditions. Integrating spatially resolved transcriptomics and proteomics allows direct mapping of gene expression and protein distribution, providing richer context for interpreting molecular and cellular organization. Moreover, methods such as MERFISH^1^, seqFISH+ ^2^, and Slide-seq^3^ have made it possible to chart high-resolution transcriptomic profiles across sections, revealing both spatial continuity and local variability. Additionally, spatial proteomics platforms such as CODEX^3,4^ and imaging mass cytometry (IMC)^3–5^ allow multiplexed protein quantification across slices, uncovering patterns of tissue architecture and function. Recent advances in spatial multiomics now enable the joint measurement of chromatin accessibility and gene expression within intact tissue architecture^6^, as exemplified by spatial ATAC–RNA sequencing and emerging tri-omic platforms, thereby enabling direct mapping of gene regulatory programs in situ. Collectively, these spatial omics approaches have transformed our capacity to interrogate complex tissue organization and heterogeneous cellular responses across diverse biological contexts.

Understanding how cells respond to perturbations within their spatial microenvironment is essential for decoding the principles that govern development, disease progression, and therapeutic response. Spatially resolved studies have uncovered how cellular behaviors and molecular programs vary dramatically depending on their position within a tissue. For instance, spatial omics revealed gene expression dynamics underlying brain development and aging^7,8^ as well as microenvironment-specific responses that drive tumor heterogeneity and influence treatment outcomes^9–12^. Yet, analyzing spatial responses to perturbations remains challenging due to the intricate interplay between cell types, local environment and global tissue organization. Even genetically identical cell types can display remarkably distinct responses depending on their spatial niche^8^, underscoring the need for computational frameworks that can resolve such context-dependent heterogeneity.

Despite the rapid advances of spatial omics, most computational frameworks remain limited in effectively integrating spatial context or employing contrastive analysis^13,14^ to capture perturbation-specific responses. Conventional methods often rely on clustering or factor decomposition without explicitly modeling spatial relationships or contrasting conditions, limiting their ability to discern subtle yet meaningful spatial differences. For instance, CellCharter^15^ models cell type distributions and neighborhood interactions but lacks the capacity for contrastive learning across conditions. Conversely, contrastive variational inference (contrastive VI)^16^ excels at disentangling shared and condition-specific latent factors but ignores spatial information, treating each observation as independent. This gap between spatial modeling and contrastive learning hinders accurate interpretation of perturbation-driven heterogeneity. Furthermore, existing frameworks have been developed exclusively for single-modality inputs, leaving the joint modeling of co-profiled chromatin accessibility and gene expression, increasingly available through spatial multiomics platforms, largely unexplored in the contrastive setting.

A natural alternative to a unified framework would be to apply contrastive and spatial methods sequentially, for instance, running ContrastiveVI to obtain condition-specific embeddings, then applying CellCharter to impose spatial structure. However, such sequential application fails to capture the mutual dependencies between spatial context and condition-specific variation. Because ContrastiveVI operates on spatially unaware embeddings, spatial covariance information is not incorporated during the disentanglement step, and post-hoc spatial smoothing cannot recover what was lost during encoding. A framework that jointly incorporates spatial niche structure within the contrastive inference process itself is therefore necessary to faithfully separate perturbation-driven signals from conserved tissue architecture.

To overcome these limitations, we developed **Haruka**, a spatially aware contrastive learning framework that disentangles condition-specific perturbations from shared tissue architecture. Haruka combines contrastive variational inference^13,16^ with an auxiliary microenvironment reconstruction task, enabling learned embeddings to capture both spatial dependencies and global condition effects. These embeddings are then aggregated into higher-order niche representations, linking fine-grained cellular responses to the broader organization of tissue microenvironments. Through this integration, Haruka establishes a generalizable framework for analyzing spatial perturbations across tissues, conditions, and omics modalities, including native support for multimodal spatial input, providing new opportunities to uncover how developmental, pathological, and therapeutic processes remodel the cellular landscape at both transcriptional and epigenetic resolution.

## Results

### Benchmarking Haruka against state-of-the-art methods reveals superior detection of salient spatial domains

Haruka is a deep generative model that jointly encodes spatial context and contrastive structure to learn two embeddings per cell: a salient (condition-specific) and a background (shared) representation. Specifically, we integrate a multi-task reconstruction objective, including spatial omics profile recovery and spatial covariance matrix reconstruction, with an additive contrastive generative process to infer both background and salient representations. These embeddings are aggregated through spatial neighborhoods and clustered to define spatial domains that capture both shared architecture and condition-specific variation (**Fig. 1**). The learned representations also support downstream analysis across multiple slices and conditions.

**Figure 1.**
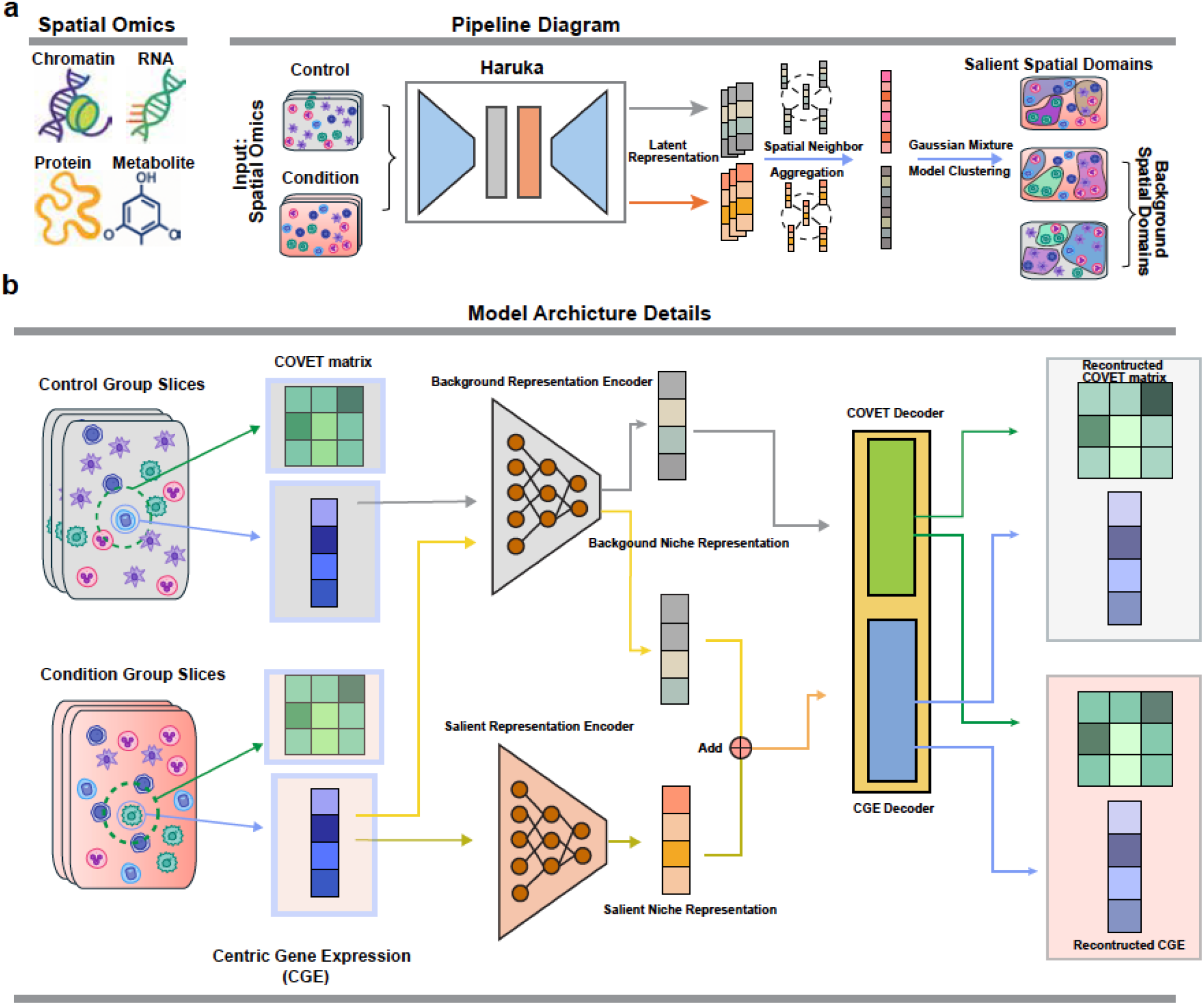
Overview of Haruka. **a**, Overview of the Haruka pipeline. Haruka takes as input multi-slice spatial omics datasets from both condition and control samples and can flexibly accommodate different spatial omics modalities. The Haruka model extracts spatial-aware salient and background representations for each niche (a centric cell with its given neighbors) after reconstruction training. The salient and background niche embeddings are then aggregated by spatial neighbors to obtain a spatially smooth niche embedding that considers multi-scale resolution information. Finally, Gaussian Mixture Model clustering is applied to obtain the final salient/background spatial domains, consistent across both condition and control slices. **b**, Details of the Haruka model architecture. Haruka first transforms each niche into two input parts: the Covariance Gene Expression matrix (COVET) of neighboring cells and the centric cell gene expression (CGE). Haruka then receives only CGE as input for subsequent encoders. For condition group niches, they are modeled by a combination of salient latent embedding and latent background embedding. Control group niches are modeled by only the background embedding. Haruka utilizes two separate decoders (CGE reconstruction layers and COVET reconstruction layers) to reconstruct CGE and COVET separately. This process forces the latent embedding to capture information contained in the niche neighbors by reconstructing COVET without explicit input. The final latent embedding (background for control groups and salient + background for condition groups) for both groups is then fed to both decoders. The model is optimized based on a variational inference framework to achieve minimal reconstruction loss in both COVET and CGE.

In simulated benchmarking, we used a MERFISH mouse hypothalamus dataset as the background and introduced region-specific perturbations to define salient domains (**Fig. 2a, Methods**). We then compared Haruka to CellCharter^15^, Contrastive VI^16^, and CINEMA-OT^17^, using ARI, NMI and related clustering metrics (**Methods**). Across tasks of varying difficulty, Haruka outperformed all other methods in detecting salient domains while maintaining competitive accuracy in background domain identification (**Fig. 2b-c and Extended Data Fig. 1**). CellCharter, which was designed primarily to capture background spatial structure, underperformed on salient variation while Haruka efficiently identified both salient and background patterns.

**Figure 2.**
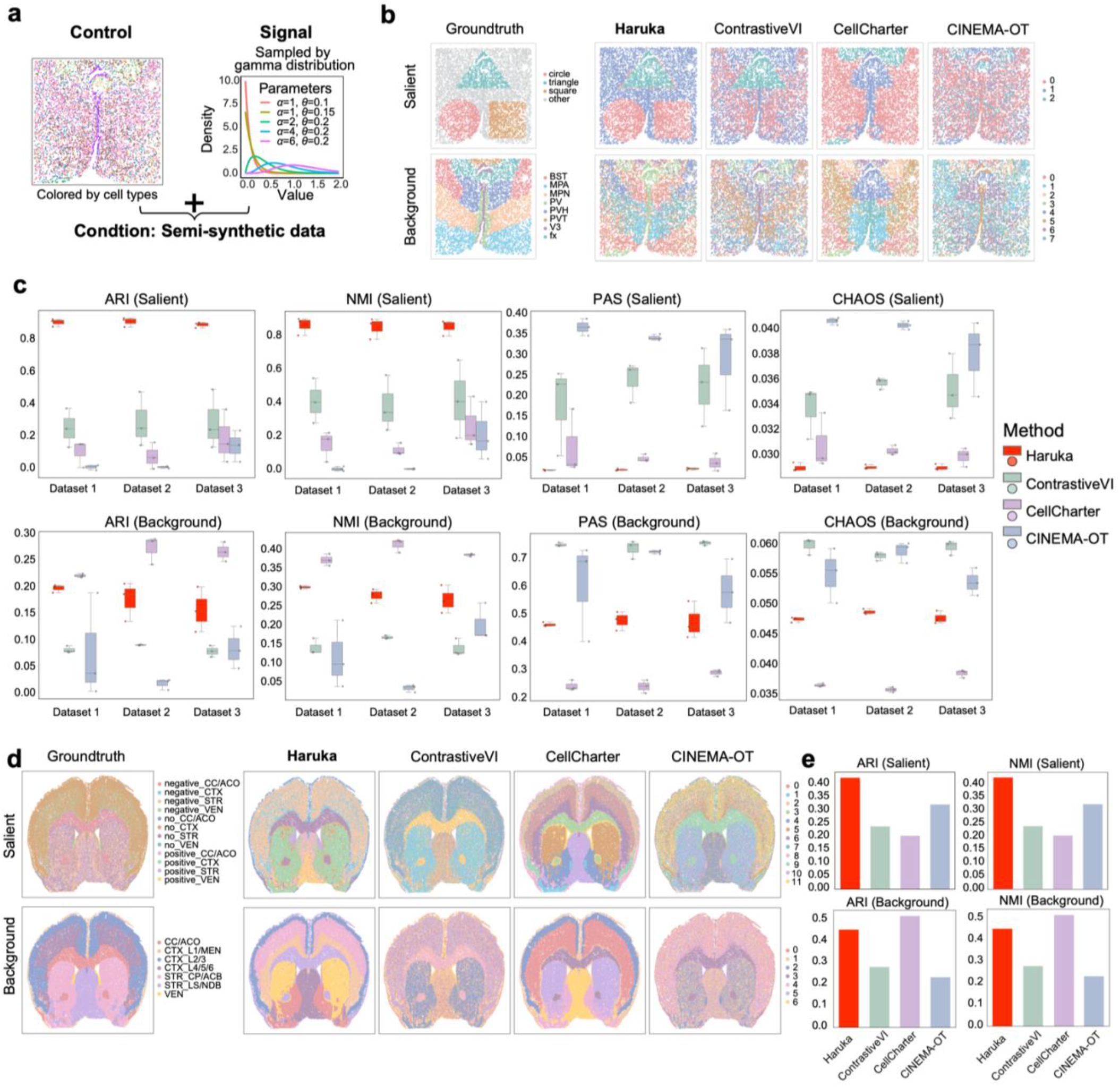
Benchmarking Haruka against existing methods using semi-simulated and mouse brain aging datasets. **a**, MERFISH mouse brain datasets simulation diagram. We add different parameter gamma distributions in three simulated spatial domains and then introduce heterogeneity shifts in these spatial domains relative to the original ground truth expression. Thus, the original brain section serves as the ground truth background spatial domain, and the three simulated spatial domain shifts represent the ground truth salient domains. **b**, Visualization of the ground truth salient and background spatial domains, and the results from Haruka alongside comparison methods (ContrastiveVI, CellCharter, and CINEMA-OT). **c**, Quantitative comparison between Haruka and comparison methods on the MERFISH semi-simulation dataset in terms of clustering accuracy (ARI, NMI) and spatial continuity (PAS, CHAOS) across different simulated salient shifts (Poisson signal parameter (alpha=2,4,6) and domain area proportion (0.1, 0.15, 0.2)). **d**, Visualization of the real-world mouse brain aging benchmark dataset. We separate each brain subregion into positively correlated, negatively correlated, and non-correlated domains with the aging procedure, totaling 3*4 = 12 salient spatial domains, with the original brain subregion serving as the ground truth background spatial domain (left). Results of Haruka and comparison methods (Contrastive VI, CellCharter, and CINEMA-OT) are also shown. **e**, Quantitative comparison of Haruka with comparison methods using clustering accuracy metrics (ARI, NMI).

For real-world evaluation, we applied Haruka to a MERFISH-based mouse brain aging dataset^8^ (**Methods**). The original study reported that gene expression responses to aging vary markedly across brain regions in a cell type-specific manner, with some regions exhibiting pronounced changes while others remained relatively stable. We used these observations to construct twelve spatial domains (four brain sections and three correlation groups per section) representing different regional responses to aging (**Methods)**. This benchmark presented two key challenges: salient domains in real tissue can be spatially discontinuous, and identical cell types may behave differently across brain sections. Because of this complexity, we excluded continuity-sensitive metrics such as CHAOS and PAS that penalize fragmented but biologically meaningful domains. Background domains were defined by anatomical subregions, and the youngest (E3.4) and oldest (E34.5) sections were designed as control and condition samples, respectively. Haruka significantly outperformed all baseline methods in identifying salient domains and performed comparably to CellCharter on background domains task (**Fig. 2d-e**). These results demonstrate that Haruka generalizes effectively across simulated and real-world datasets, capturing biologically relevant heterogeneity with robustness and scalability.

### Haruka identifies salient immune domains associated with immunotherapy response in melanoma

To assess Haruka’s performance in a clinical context, we applied it to a large CODEX melanoma spatial proteomics dataset^9^ analyzed before and after immune checkpoint blockade (ICB) therapy and annotated with clinical status as a responder or non-responder to treatment based on objective response rate (ORR) labels (n = 6 patients total). In our analysis, pre-treatment tissues were designated as control, and post-treatment tissues as condition, allowing Haruka to identify response-associated spatial variation (**Fig. 3a**). Haruka discovered four recurrent salient immune domains (Domains 0-3) that capture microregional heterogeneity in post-treatment tumor landscapes (**Fig. 3b-e and Extended Data Fig. 2a**).

**Figure 3.**
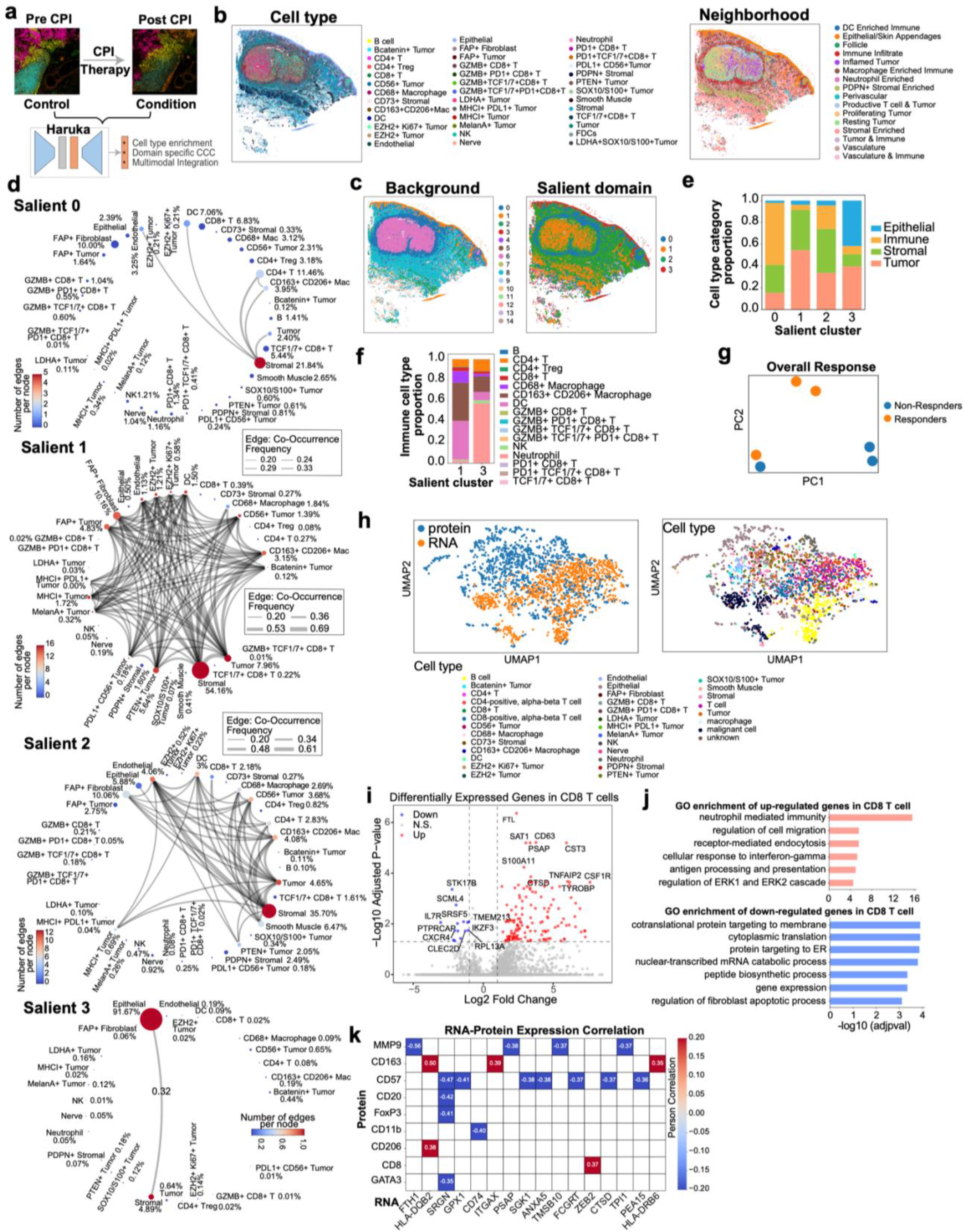
Haruka-derived microenvironment reveals immunotherapy response heterogeneity in the CODEX melanoma dataset. **a**, Overview of the Haruka pipeline applied to the melanoma immunotherapy CODEX dataset. **b**, Spatial distribution of annotated cell types and cell neighborhoods in a representative tissue section. **c**, Background and Salient spatial domains derived from Haruka in the same representative tissue section as b. **d**, Cell type proportion and cell-cell co-occurrence for each salient spatial domain (0,1,2,3). Spot size indicates cell type proportion in this domain, and edge abundance between two nodes denotes the cell type co-occurrence frequency in this domain. Four salient spatial domains show distinguished cell type and co-occurrence patterns, reflecting their heterogeneous response to immunotherapy. **e**, Coarse-grained cell type proportion comparison among four salient spatial domains. **f**, Fine-grained immune cell type proportion comparison between salient spatial domains 1 and 3, which show closer coarse-grained immune cell type proportion distribution. **g**, Principal Component Analysis (PCA) results for salient spatial domain frequency for each patient with Non-responder/Responder phenotype annotation. **h**, UMAP visualization of MaxFuse co-embedding of external scRNA-seq dataset and salient domain 0 CODEX proteomics data. **i**, Volcano plot showing differential gene expression results in CD8 T cells from scRNA-seq data grouped by matched/non-matched CODEX salient domain 0. **j**, GO term enrichment results on the significant differential expression marker genes in matched domain 0 cells (resistant to immunotherapy). **k**, RNA-protein co-expression correlation in domain 0 with the integrated result after MaxFuse. Higher PCC indicates potential higher co-expression of the spatial proteomics biomarker and gene expression.

Importantly, Haruka revealed that patients with objective responses (ORR=1) showed enrichment for Domains 0 and 2 in this dataset, each representing distinct yet converging routes to effective ICB response (**Extended Data Fig. 2b**). Both domains were characterized by networks centered on TCF1^+^ CD8^+^ T cells, a stem-like precursor subset essential for sustained antitumor immunity^18,19^. In concert with dendritic cells (DCs) and CD4^+^ T cells^20^, these form a productive immune circuit necessary for T cell priming and expansion. Domain 0 represented a T cell dominated activation hub (**Fig. 3d**), where TCF1^+^ CD8^+^ T cells directly interacted with tumor cells^21^, while Domain 2 formed a stromal-integrated immune zone where stromal cells supported rather than impeded immune–tumor interactions. Here, TCF1^+^ CD8^+^ T cells remain central, but function within a broader, more balanced co-occurrence structure that maintains immune accessibility^19,20^. Thus, both domains represent effective immunotherapy microenvironments: one via immune density, the other via immune-stromal harmony.

In contrast, patients without objective responses (ORR = 0) showed enrichment for Domains 1 and 3 in this dataset (**Extended Data Fig. 2b**), two distinct modes of immunotherapy resistance. A shared hallmark is a near-complete absence of TCF1^+^ CD8^+^ T cells, a progenitor population critical for sustaining anti-tumor immunity and predictive of checkpoint blockade responsiveness^18^. Domain 1 was dominated by interconnected fibroblasts and M2-like macrophages that form a non-productive, immunosuppressive niche (**Fig. 3d**), consistent with immune exclusion mechanisms^22,23^. Domain 3 displayed an epithelial-dominant phenotype lacking immune cell infiltration (**Fig. 3d-e**), consistent with immune desert^24^. The absence of dendritic cells, CD4^+^ T cells, and cytotoxic CD8^+^ subsets (**Fig. 3f**) suggest a structurally inert, antigen-silent environment resembling that of so-called “cold tumors”^25,26^. Thus, both Domains 1 and 3 exhibit resistance modes driven by spatial exclusion and immune invisibility.

Taken together, our findings support that the distribution of salient spatial domains captures biologically meaningful differences between patients with and without objective responses, consistent with established immune correlates of checkpoint blockade efficacy. Principal component analysis (PCA) of salient domain fractions separated patients according to ORR in this dataset (**Fig. 3g**), concordant with the known association between TCF1^+^ CD8^+^ T cell infiltration and checkpoint blockade response. We note that these findings are derived from a small cohort (n = 6 patients) and should be considered hypothesis-generating; prospective validation in larger, independent cohorts will be required to assess clinical utility. Importantly, Haruka uses only simple, binary labels (pre-vs. post-treatment), highlighting its flexibility and applicability in clinical research settings.

To further explore the biological basis of these salient domains, we integrated the ORR-negative Domain 3 CODEX reference with an independent post-immunotherapy melanoma scRNA-seq datasets^27^ using MaxFuse^28^ (**Fig. 3h and Methods**). In differential gene expression analysis of matched CD8 T cells, we identified transcriptional signatures characterized by inflammatory activation but global translational repression. These included downregulation of biosynthetic and ER-targeting machinery and enrichment of interferon and antigen presentation pathways **(Fig. 3i-j and Methods)**, consistent with known immune exclusion mechanisms^29–32^. These changes suggest an immune exhaustion driven by chronic cytokine exposure and TGF-β signaling, consistent with fibroblast-mediated immune suppression in the TME (tumor microenvironment)^33–36^ and cancer-associated fibroblast (CAF) barriers that restrict immune cell infiltration^37,38^. In RNA-protein correlation analysis (**Fig. 3k**), we further showed that immunosuppressive markers CD163 and CD206 were correlated with tolerogenic transcripts such as SIGLEC10 and SPON1^39–41^, defining an M2-like macrophage program associated with checkpoint therapy failure.

Together, these findings define a spatially encoded model of immune resistance in melanoma where immune cells persist but exhibit translational suppression and metabolic exhaustion.

### Haruka captures salient and background representations to map fibrosis progression in human lung

To demonstrate Haruka’s flexibility across spatial omics platforms, we applied Haruka to a large-scale Xenium dataset of human pulmonary lung fibrosis^42^ (**Fig. 4a and Methods**). In this setting, 10 normal human lung tissue samples were designated as the control group, while 35 lung tissues from patients with pulmonary fibrosis served as the condition group (**Fig. 4a**). Inter-sample technical variation was addressed through per-cell normalization and quality control filters applied uniformly across all samples prior to model training (**Methods**). We found that Haruka’s background and salient domains corresponded closely with transcript-based and cell-based niches defined in the original study, suggesting that Haruka can faithfully recover known tissue architecture (**Fig. 4b and Extended Data Fig. 3a-b**).

**Figure 4.**
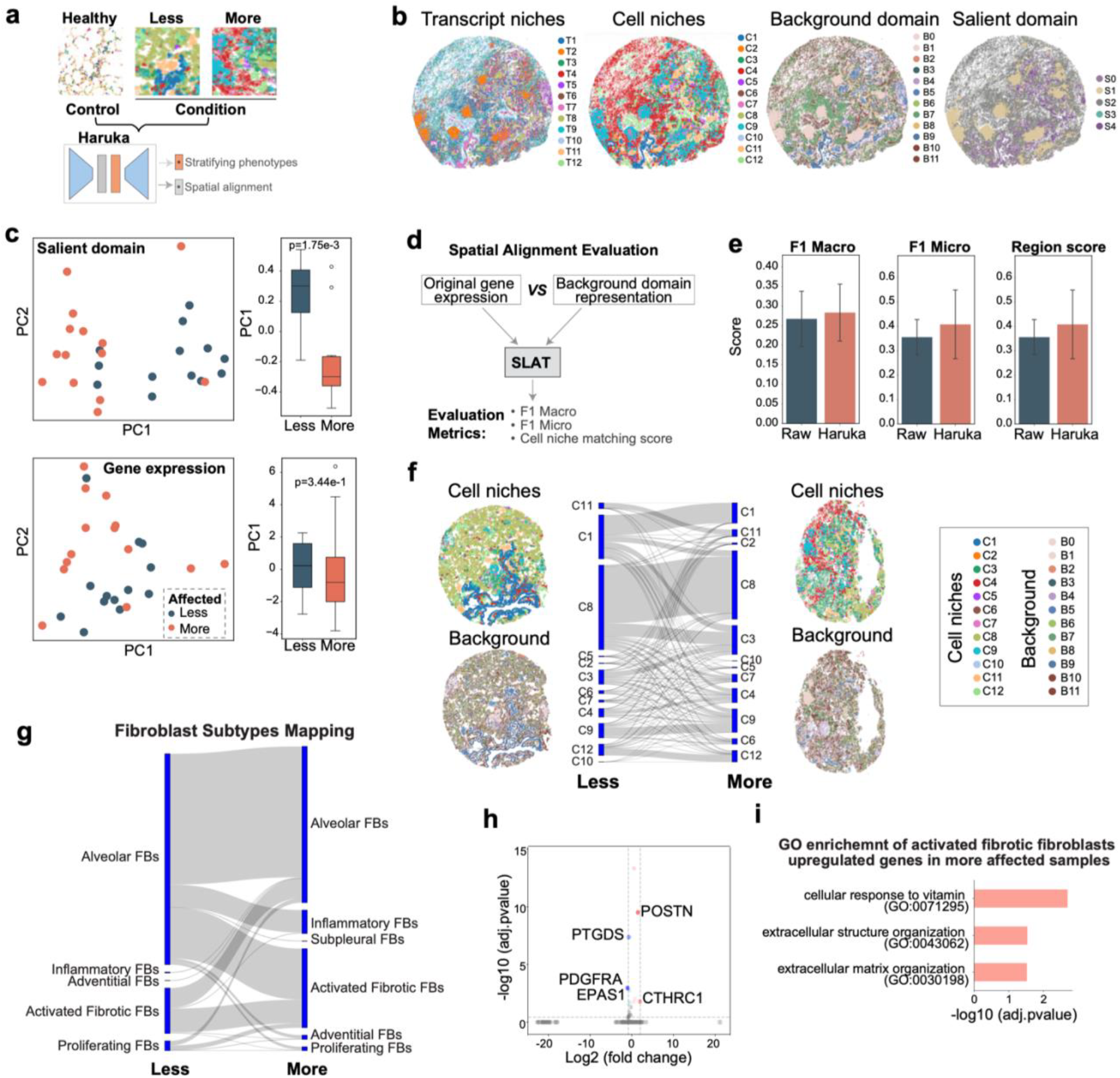
Haruka captures lung fibrosis dynamics at niche resolution. **a**, Overview of the Haruka pipeline applied to the human lung fibrosis 10x Xenium dataset. **b**, Visualization of annotated transcript and cell niches, Haruka derived background and salient domains in a representative tissue section. **c**, Comparison of separation between less and more affected samples using Haruka’s salient domain frequency vector (top) versus raw gene expression (bottom). Left: PCA visualization of samples labeled as less or more affected in the original study. Right: Boxplots of PC1 values for the two groups, with rank-sum test *p*-values shown. **d**, Diagram of our cross-condition slice alignment benchmarking. We compared the salient background representation with original gene expression, utilized as input for the SLAT alignment tool, to align the cross-condition slices (less-more affected samples) and compare the quality of each representation by the final alignment metrics (F1-macro/micro and matching accuracy regarding the annotated CN niche label). **e**, Quantitative evaluation of comparisons in the three metrics (F1-Macro, F1-micro, and Region score) across three less-more slices pairs. **f**, Matched Sankey plot for the CN niche identity in aligned less affected and more affected slice cells in the representative sections. Cell niches and background domains were also visualized for both slices. **g**, Potential cell type transition from less affected to more affected situation by the SLAT matched results based on Haruka background representation. **h**, Volcano plot for the differential expression analysis for the Alveolar FBs in the less affected samples grouped by the label (will transfer to Activated Fibrotic FBs in the future and won”t). X-axis represents the log2 transformed fold change, and y-axis represents −log10 transformed adjusted p-values. Five top genes were highlighted. **i**, Gene ontology enrichment of upregulated genes in “will transfer to Activated Fibrotic FBs Alveolar FBs” group defined in **g**.

To evaluate whether salient domains could capture fibrosis progression, we projected samples into low-dimensional spaces derived from Haruka’s salient representations and from raw gene expression (**Methods**). We found projection of samples using Haruka’s salient embeddings separated mildly affected versus severely fibrotic tissues, a distinction that raw gene expression failed to capture (**Fig. 4c**). These results demonstrate that Haruka’s salient domains provide a more sensitive representation of disease progression, overcoming the confounding effects of spatial complexity and intra-group variation.

We next tested whether Haruka’s background representations could improve tissue slice alignment across conditions. Using the SLAT algoritm^43^, we compared alignments based on original gene expression profiles versus Haruka-derived background embeddings across fibrosis severity (**Fig. 4d and Methods**). Haruka substantially improved CNiche-matching and F1 scores (**Fig. 4e**) indicating more accurate mapping between control and fibrotic slices. This alignment revealed transitions from alveolar fibroblasts (FBs) to inflammatory fibroblasts, a hallmark of lung fibrosis pathology (**Fig. 4f-g**).

To identify early transcriptional priming events preceding fibroblast activation, we stratified alveolar FBs in mildly affected tissue according to whether their matched cells in severely affected tissue adopted a fibrotic identity (**Fig. 4g**). Differential gene expression analysis revealed upregulation of extracellular matrix organization and structure pathways in the pre-transition subgroup (**Fig. 4h-i**), consistent with early activation towards a fibrotic state. These programs also align with single-cell studies that have consistently identified ECM-producing fibroblast subsets as central to fibrotic tissue development^44–46^.

### Haruka dissects the microenvironmental basis of KRAS inhibitor resistance in lung cancer

To assess the translational applicability of Haruka, we employed a KRAS^G12D^-mutant mouse model of metastatic lung cancer and examined the therapeutic effects of a KRAS^G12D^ specific inhibitor (MRTX1133) using spatial transcriptomics (**Fig. 5a**). Using the 10x Genomics Xenium platform, we generated high-resolution spatial maps of whole lung specimens that contained disseminated KRAS^G12D^-mutant lung tumors. We compared specimens that were treated with vehicle control versus those subjected to MRTX1133 treatment (n = 4 specimens per treatment condition) (**Methods and Extended Data Fig. 4a-d**).

**Figure 5.**
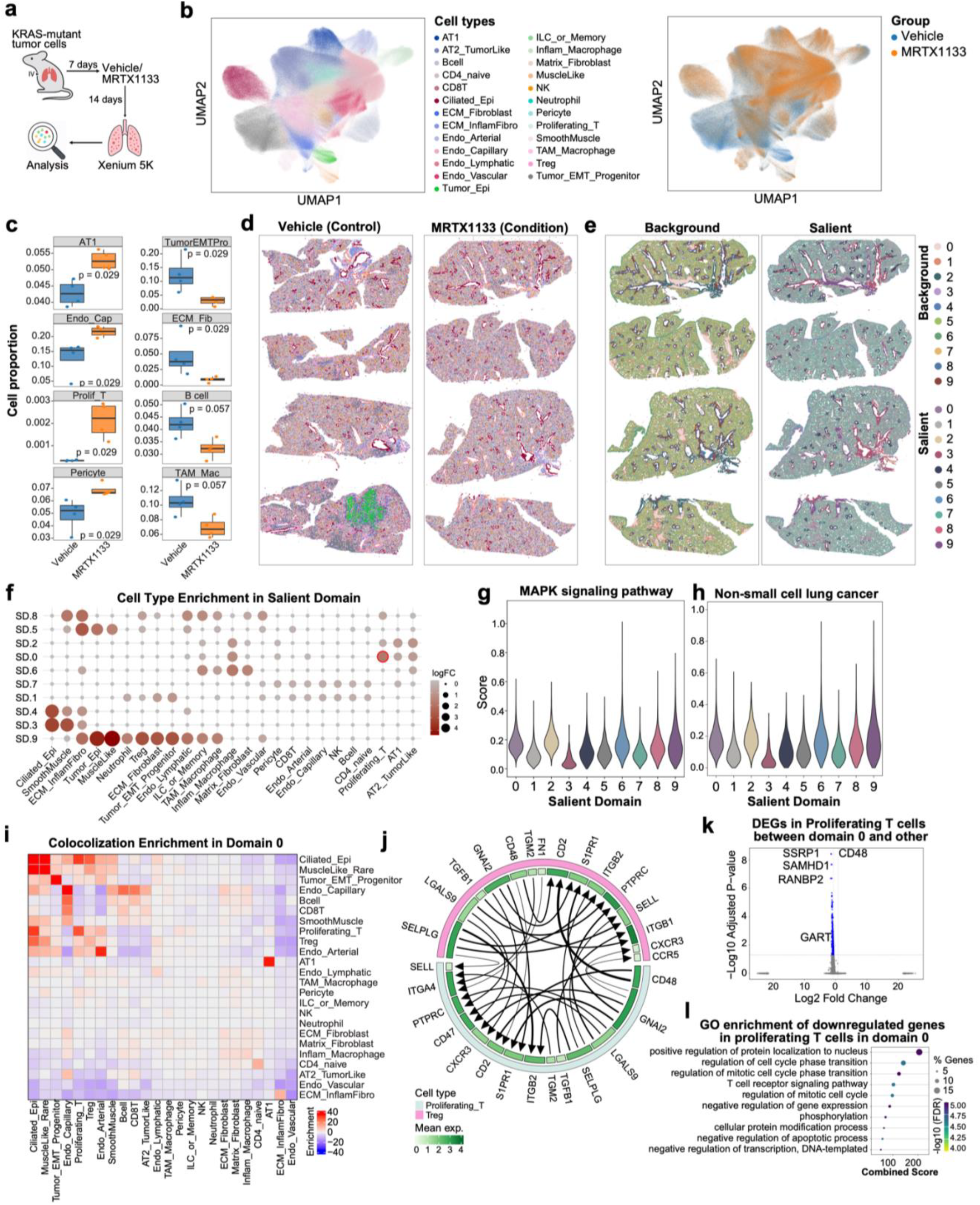
Haruka dissects the microenvironmental basis of KRAS inhibitor resistance in lung cancer. **a**, Experimental design for generating the Xenium 5K dataset. KRAS^G12D^-mutant tumor cells were intravenously injected into mice and treated with either vehicle or the KRAS^G12D^ inhibitor MRTX1133 for 5 days, followed by 9-day washout and spatial transcriptomic profiling on day 21. **b**, UMAP visualization of cell type annotations across all samples, highlighting both the cell type identities (left) and treatment groups (right). **c**, Comparison of cell type proportions between Vehicle and MRTX1133-treated tumors (n = 4 biological replicates per group). *P*-values were calculated using the Wilcoxon rank-sum test. Treatment with MRTX1133 increases the proportion of epithelial cells (AT1) while decreasing immune cell fractions, including B cells and tumor-associated macrophages (TAM_Mac). **d**, Spatial maps showing cell type distributions across biological replicates for Vehicle and MRTX1133-treated tumors. Colors correspond to cell types shown in panel b. **e**, Spatial visualization of Haruka-derived background and salient domains across four MRTX1133-treated replicates, showing distinct domain-specific spatial organization. **f**, Cell type enrichment across salient domains, with dot size and color representing log-transformed fold change of enrichment significance. **g-h**, Distribution of pathway activity scores for the MAPK signaling and non–small-cell lung cancer pathways across salient domains, highlighting differential pathway activation patterns. **i**, Domain-specific colocalization enrichment map showing cell-cell proximity patterns in salient domain 0; higher enrichment values indicate stronger colocalization between corresponding cell-type pairs. **j**, Cell-cell communication network in salient domain 0 between proliferating T cells and regulatory T cells (Tregs), where arrows indicate ligand-receptor signaling direction. The outer ring color denotes the interacting cell types, while the inner ring color reflects the expression level of the corresponding ligand or receptor genes. **k**, Differential gene expression analysis of proliferating T cells in domain 0 compared with all other domains, revealing distinct transcriptional programs associated with resistant niches. The top five differentially expressed genes are highlighted. **l**, Gene Ontology (GO) enrichment analysis of downregulated genes in proliferating T cells within domain 0, identifying pathways linked to transcriptional repression, cell-cycle regulation, and phosphoregulation.

We identified diverse epithelial, stromal, and immune populations, including tumor-like alveolar type 2 (AT2) cells, fibroblast subtypes, and multiple immune lineages (**Fig. 5b**). Stratification by treatment revealed broadly overlapping cellular distributions between vehicle- and MRTX1133-treated tumors, indicating consistent capture of major cell types across conditions (**Fig. 5b and Extended Data Fig. 4e**). We next quantified treatment-associated shifts in cellular composition. MRTX1133-treated tumors exhibited a marked expansion of AT1 cells, endothelial capillaries, proliferating T cells, and pericytes, alongside a significant depletion of tumor EMT progenitors and ECM fibroblasts (**Fig. 5c and Extended Data Fig. 4f**). We also observed a trend toward reduced B cells and tumor-associated macrophages, although these changes did not reach statistical significance. Notably, discrete regions enriched for KRAS/TP53 (KP) tumor epithelium were observed, consistent with the TP53-deficient subtype (**Fig. 5d**).

Using MRTX1133-treated samples as controls and vehicle-treated samples as conditions, Haruka identified stable background domains alongside distinct salient domains that capture localized intra-condition heterogeneity (**Extended Data Fig. 4g–h**). These patterns indicate that Haruka resolves biologically meaningful spatial variation rather than technical noise, including the focal emergence of KP tumor epithelium. In particular, salient domains 1 and 5 were enriched for inflammatory macrophages and tumor-associated macrophages (**Extended Data Fig. 4i–j**), suggesting domain-specific immune remodeling within the tumor microenvironment.

To further demonstrate Haruka’s versatility, we next reframed the analysis to investigate mechanisms of KRAS inhibitor (KRASi) resistance. Specifically, we designated KRAS inhibitor–treated tumors as the condition and vehicle-treated tumors as the control, enabling identification of spatial features associated with reduced drug response. This analysis focuses on how spatially complex microenvironments shape treatment outcomes, even within the same cell type (**Fig. 5d-f, Extended Data Fig. 5a and Methods**).

Because effective KRAS inhibition should suppress downstream MAPK signaling, we reasoned that elevated pathway activity would signify a reduced drug response. Comparative analysis revealed that salient domains 0, 2 and 6 maintained higher MAPK and lung cancer pathway activity than other domains, identifying them as candidate resistance-associated niches within the tumor microenvironment (**Fig. 5g-h**). When we examined the TME of each domain by comparing spatial colocalization enrichment against all other domains (**Fig. 5i and Extended Data Fig. 5b**), we found that Domain 0 exhibited strong colocalization of tumor epithelial cells with tumor EMT progenitors, ciliated epithelial cells, and proliferating T cells (uniquely enriched in domain 0, as **Fig. 5f**), suggesting an active proliferative and transitional niche (**Fig. 5i**). We also observed enrichment of Tregs in close proximity to tumor and stromal elements and enhanced interactions consistent with angiogenic remodeling.

To characterize the immunoregulatory network of domain 0, we performed ligand-receptor analysis focused on proliferating T cells and Tregs (**Methods**). We revealed a dense adhesion- and trafficking-centered signaling module to support an immunosuppressive microenvironment (**Fig. 5j**). We found that Tregs contributed key ligands, including LGALS9, SELPLG, and TGFB1, while proliferating T cells expressed complementary receptors such as ITGB1/2, SELL, S1PR1, and CD2. Notably, integrin-mediated activation of latent TGF-β positions Tregs as a central source of TGF-β-driven immunosuppressive signaling^22^. These paired interactions form a bidirectional integrin-selectin and CD2-CD48 axis, coordinating T-cell retention, positioning, and contact-dependent regulation. Together, this creates a dense adhesion-migration-regulation circuit that stabilizes a spatially cohesive, TGF-β–driven immunosuppressive microenvironment.

To further dissect the transcriptional state of proliferating T cells in domain 0, we identified differentially expressed genes relative to all other domains. Many were downregulated, including key regulators of DNA replication and cell cycle progression (SSRP1^47^, SAMHD1^48^, and RANBP2^49^, **Fig. 5k**). Gene ontology enrichment analysis revealed broad suppression of T cell receptor signaling, cell cycle transition, protein phosphorylation, and regulation of apoptotic processes (**Fig. 5l**). Despite their spatial abundance, proliferating T cells exhibit transcriptional repression of proliferative and effector programs, consistent with functional exhaustion and immune suppression within the resistant tumor microenvironment^50^.

Together, these results show that candidate KRASi resistance may be spatially organized through microenvironments that couple proliferative tumor programs and vascular remodeling with coordinated Treg-mediated immunosuppression, thereby limiting T cell function. Haruka identifies these candidate resistance-associated niches and their signaling architecture, providing a framework for uncovering therapeutic vulnerabilities within complex tumor ecosystems.

### Haruka resolves multimodal regulatory heterogeneity in spatially co-profiled chromatin and transcriptomic landscapes during neuroinflammation

To assess Haruka’s capability in the spatial multimodal setting, we applied it to spatially resolved multiomics data co-profiling RNA expression and ATAC chromatin accessibility from a P21 mouse brain under lysolecithin (LPC)-induced neuroinflammatory perturbation, generated using ARCseq — a spatial platform that captures transcriptomic and epigenomic profiles at matched spatial coordinates within intact tissue sections^51^ (**Fig. 6a**). As a proof-of-concept application using a single pair of matched tissue sections, spatial multiomics slices from an uninjured P21 mouse (P21S3) were designated as control, and paired slices from an LPC-injected P21 mouse (LPC21S1) were designated as condition. LPC injection is a well-established model of focal demyelination and neuroinflammation, as lysolecithin disrupts myelin membranes and induces spatially heterogeneous inflammatory and repair responses^51^.

**Figure 6.**
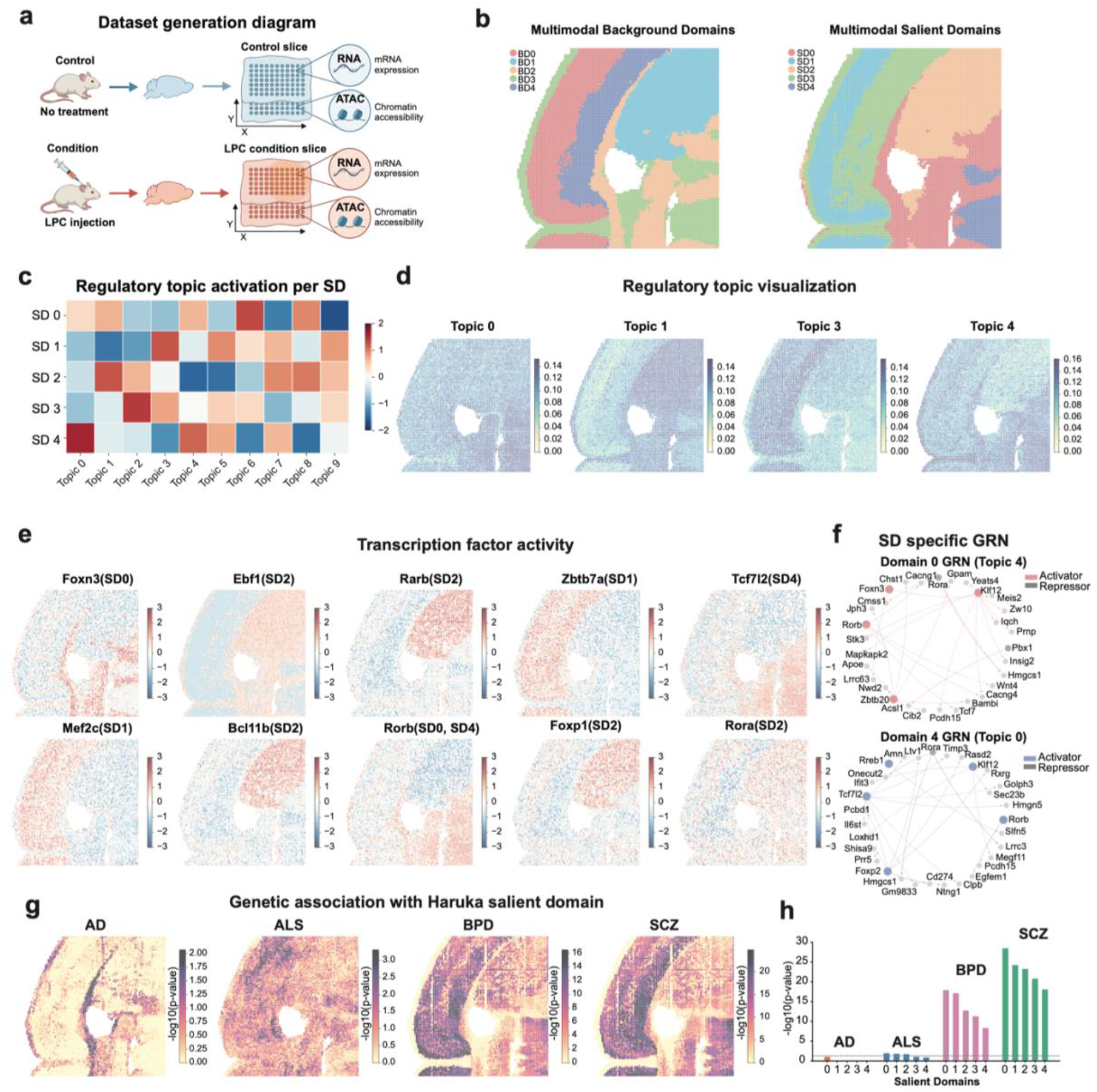
Multimodal spatial domain identification and gene regulatory network inference in LPC-induced demyelination. **a**, Schematic of the experimental design. Spatial ATAC–RNA-seq (ARCseq) was performed on a control (P21, no treatment) and a condition (P21, LPC injection) mouse brain section, generating matched chromatin accessibility and gene expression profiles per spot. **b**, Spatial maps of Haruka multimodal background domains (left) and salient domains (right), identified by applying ContrastiveVI independently to RNA and ATAC, followed by CellCharter spatial aggregation, concatenation, and clustering (k = 5). **c**, Heatmap of mean scDoRI regulatory topic activation (z-scored) across the five salient domains. Topics 0, 1, 3, and 4 show domain-specific activation. **d**, Spatial visualization of scDoRI topic weights for Topics 0, 1, 3, and 4. **e**, Spatial maps of TF activity scores for ten selected TFs, annotated by their primary domain enrichment. **f**, Domain-specific gene regulatory networks for Domain 0 (Topic 4, top) and Domain 4 (Topic 0, bottom). Nodes represent TFs (large) and target genes (small); edges indicate activating (colored) or repressing (gray) relationships (permutation test, 1,000 permutations, p < 0.05). **g**, Spatial maps of gsMAP −log_10_(p) for AD, ALS, BPD, and SCZ. (h) Cauchy combination test results (−log_10_(p_Cauchy)) per salient domain for all four traits.

Haruka was independently trained on each modality then aggregated into a joint multimodal embedding by concatenating the spatially smoothed salient and background latent representations from both modalities (**Methods**). This modality-parallel design enables Haruka to disentangle perturbation-induced chromatin remodeling from shared tissue architecture, independently within each layer, before integrating them into a unified niche representation. Haruka decomposed the data into five background domains capturing shared spatial tissue structure and five salient domains (SDs) representing condition-specific variation (**Fig. 6b**). Background domains recovered major anatomical subregions including cortical layers, white matter, and subcortical structures, while salient domains identified spatially localized LPC response patterns distributed across the cortex, deep brain regions, and the lesion-proximal zone.

To characterize the regulatory programs underlying each salient domain, we applied scDoRI^52^, a spatial multiomics topic model that jointly decomposes ATAC accessibility, transcription factor (TF) expression, and RNA expression into interpretable regulatory topics, and subsequently infers domain-specific gene regulatory networks (GRNs) (**Methods**). scDoRI identified ten regulatory topics, of which Topics 0, 1, 3, and 4 showed the most pronounced differential activation across salient domains (**Fig. 6c,d and Extended Data Fig. 6**). SD4 was primarily driven by Topic 0, and SD0 by Topic 4, while SD2 and SD1 exhibited enrichment for Topics 1 and 3, respectively; SD3 showed diffuse, low-level activation consistent with a sparse or transitional state (**Fig. 6c**). Spatial visualization confirmed that these four topics mark anatomically coherent zones — Topic 0 concentrating in thalamocortical territories, Topic 1 in projection neuron corridors, Topic 3 in deep excitatory layers, and Topic 4 in upper cortical and lesion-proximal zones (**Fig. 6d**).

Domain-specific transcription factor activity analysis revealed pronounced differences in regulatory identities across salient domains (**Fig. 6e**). SD0 was characterized by Foxn3 and Rora activity, consistent with an upper/mid-cortical and LPC-perturbed astrocytic state. SD4 exhibited elevated activity of Tcf7l2 and Rorb, consistent with a thalamocortical identity in which Rorb defines layer 4 transcriptional programs and Tcf7l2 marks Wnt-responsive neuronal progenitors. SD2 was enriched for Ebf1, Rarb, Bcl11b, and Foxp1 — regulators associated with medium spiny neurons and deep-layer projection identity and remyelination programs. SD1 showed Zbtb7a and Mef2c activity, consistent with activity-dependent deep-layer excitatory neuron programs. The shared presence of Rorb across SD0 and SD4 suggests a convergent accessibility state at cortical-thalamic boundaries despite divergent perturbation responses.

By integrating gene expression and chromatin accessibility, we next inferred domain-specific GRNs to characterize the underlying regulatory mechanisms (**Fig. 6f**). SD0 (mapped to Topic 4) was dominated by a regulatory program centered on Foxn3 and Rora as upstream activators, converging on target genes enriched for lipid metabolic and neuroinflammatory response pathways — including Hmgcs1 and Insig2 (cholesterol biosynthesis), Acsl1 (long-chain fatty acid metabolism), and Apoe (neuroinflammatory lipid transport). This is consistent with prior studies demonstrating that cholesterol biosynthesis is required for remyelination and that APOE plays a central role in neuroinflammatory responses^53,54^. Additional repressor connections from Zbtb20 and Klf12 further shaped the SD0 transcriptional landscape, marking this domain as a perturbation-induced metabolic-inflammatory regulatory state. In contrast, SD4 (mapped to Topic 0) was governed by Rorb, Rora, and Foxp2 as activators, with Tcf7l2 and Klf12 as additional regulators, whose target programs included synaptic organization, axonal guidance, and circuit identity genes — consistent with Rorb defining cortical layer 4 transcriptional identity and Foxp2 marking deep-layer projection neurons^55,56^. Together, these findings indicate that Haruka resolves LPC-induced neuroinflammatory heterogeneity into distinct mechanistic regulatory states: SD0 as a perturbation-induced lipid-metabolic and inflammatory response, and SD4 as a resilient neuronal regulatory architecture.

To assess the disease relevance of these salient domains, we performed gsMAP^57^-based GWAS enrichment analysis, mapping human complex trait genetic signals onto Haruka’s salient domain representations (**Fig. 6g,h**). We profiled four GWAS traits: Alzheimer’s disease (AD), amyotrophic lateral sclerosis (ALS), bipolar disorder (BPD), and schizophrenia (SCZ). At the spatial spot level, AD showed minimal enrichment (4.2% of spots at p < 0.05, no Bonferroni-significant spots), and ALS showed modest enrichment (29.0%, no Bonferroni correction survived), consistent with the limited genetic architecture shared between these neurodegenerative diseases and the cortical/thalamocortical domain programs captured here. In contrast, BPD and SCZ showed strong and spatially concentrated enrichment (78.1% and 83.3% of spots at p < 0.05; 4,149 and 5,301 Bonferroni-significant spots, respectively), with the Cauchy combination test revealing that both SD0 and SD4 drove the dominant enrichment signal (**Fig. 6h**). This pattern is consistent with large-scale genetic studies demonstrating that schizophrenia and bipolar disorder share substantial genetic architecture and that schizophrenia risk loci are enriched for neuronal and synaptic processes ^58,59^ Domain-level diagnostic gene analysis identified high Pearson correlation coefficients between Haruka salient representations and GWAS genes including CACNA2D1, NRG1, R3HDM1, and SEZ6L2 for SCZ and BPD, concentrated in Domains 1 and 4 (**Extended Data Fig. 7**).

Notably, the GRNs associated with SD0 and SD4 correspond to two complementary disease-relevant regulatory programs: SD4 reflects intrinsic neuronal regulatory architecture linked to circuit identity and synaptic organization, while SD0 captures a perturbation-induced lipid metabolic and neuroinflammatory response state. Together, these results suggest that Haruka, by jointly modeling transcriptomic and chromatin accessibility information in the spatial contrastive framework, can link spatially resolved multimodal regulatory programs with human disease genetics, providing a principled approach to understanding how neuroinflammatory heterogeneity interfaces with the genetic architecture of psychiatric disease.

## Discussion

This study presents spatial contrastive analysis, a computational framework for defining salient (condition-specific) and background (shared) spatial domains. Motivated by the rapid growth of large-scale spatial omics datasets spanning control and perturbed states, our approach addresses a central question in spatial biology: how do perturbations alter tissue architecture and what factors drive these changes.

In contrast to dissociated single-cell data, where each cell is analyzed in isolation, cellular responses within intact tissues are shaped by their spatial context. After a perturbation, a cell’s molecular profile reflects both the direct influence of the perturbation and indirect influences from neighboring cells that have themselves been altered. This interdependence makes it nearly impossible to apply conventional single-cell methods, which lack spatial information, to accurately analyze and compare perturbation-induced changes. Understanding such interactions is critical for identifying spatial niches that influence outcomes such as immunotherapy response^10^, or or the neurodegenerative processes in Alzheimer’s-related diseases^60^.

Haruka, however, reframes this problem within the spatial contrastive analysis framework: it separates potential noisy variations induced by neighboring cells (but not by the perturbation itself). This approach forces the model to disentangle the salient part from the background, thereby extracting the changes introduced by the perturbation. Moreover, this approach can reveal perturbation-induced biological processes that are unidentifiable by previous methods. For instance, Haruka can help biologists capture subtle but crucial changes, such as a cell type without significant gene expression but with a completely different microenvironment, a common feature in neurodegeneration or aging processes^8,60^.

Across the four biological applications presented here — melanoma immunotherapy response, pulmonary fibrosis progression, KRAS inhibitor response in lung cancer, and LPC-induced neuroinflammation — a unifying principle emerges: the spatial microenvironment is not merely a backdrop for cellular behavior but an active determinant of how cells interpret and respond to perturbations. In each context, cells of the same type displayed markedly different molecular programs depending on their niche identity, and these niche-level differences carried biological and clinical significance that transcended cell-type-level analysis. The ability of Haruka to recover these microenvironmental distinctions, across platforms ranging from CODEX proteomics to Xenium transcriptomics to spatial multiomics, underscores the generalizability of spatial contrastive analysis as a conceptual and computational approach.

The extension of Haruka to spatially resolved multiomics data further demonstrates its flexibility as a generalizable contrastive framework. When applied to ARCseq data co-profiling RNA and ATAC from the same tissue sections, Haruka successfully disentangles condition-specific chromatin remodeling from shared tissue architecture, independently within each modality, before fusing them into a unified niche representation. This modality-parallel design preserves modality-specific signal while enabling cross-modal complementarity — for instance, identifying regulatory programs in the chromatin layer that are not apparent from transcriptome alone, such as the cholesterol biosynthesis program primed in LPC-perturbed cortical niches. Integration with scDoRI topic modeling and GRN inference further demonstrates that Haruka’s salient domain representations serve as biologically coherent anchors for downstream regulatory analysis, linking perturbation responses to transcription factor networks and human disease genetics. We anticipate that as spatial multiomics platforms continue to mature and expand to higher-order modality combinations — including spatial chromatin conformation, protein, and metabolite profiling — the modality-parallel architecture of Haruka will provide a natural framework for integrating these richer molecular landscapes in the contrastive setting.

Furthermore, with advancements in current profiling techniques, the co-profiling of gene knockout and spatial transcriptomics are already yielding promising results^61^. Haruka can be applied to such datasets to handle complex data situations and decode more information about real tissue responses after gene knockout. Additionally, extending Haruka to integrate images from H&E-stained tissues holds promise for improving its performance in clinical pathology settings. Adopting this framework for applications involving continuous conditions (e.g., development, aging, and other temporal situations) instead of binary control conditions also presents a promising avenue for the Haruka framework.

In summary, Haruka provides a unified framework for analyzing multi-slice spatial omics datasets annotated with control and condition information. By defining salient and background spatial domains, it captures both perturbation-induced remodeling and conserved tissue architecture (e.g., brain sections or basic tissue components) across conditions. These representations enable diverse downstream analyses - such as differential cell-cell interactions, niche composition, and communication networks - similar to general spatial analysis pipelines but with explicit modeling of contrastive context.

As with other deep generative models, Haruka requires GPU compute, which may limit use on CPU-only systems; scalability to very large datasets (>10M cells) will benefit from continued optimization. As an unsupervised method, domain annotation benefits from domain expertise, a step that will become more tractable as spatial cell atlases mature. Most importantly, the proof-of-concept multiomics application presented here relied on a single pair of tissue sections; future work applying Haruka across multiple spatial multiomics replicates and independent neuroinflammation datasets will be essential to establish the reproducibility of the condition-specific regulatory programs identified here, and to more firmly connect spatially resolved epigenomic heterogeneity to the genetic architecture of psychiatric disease.

## Figures

**Extended Data Figure 1.**
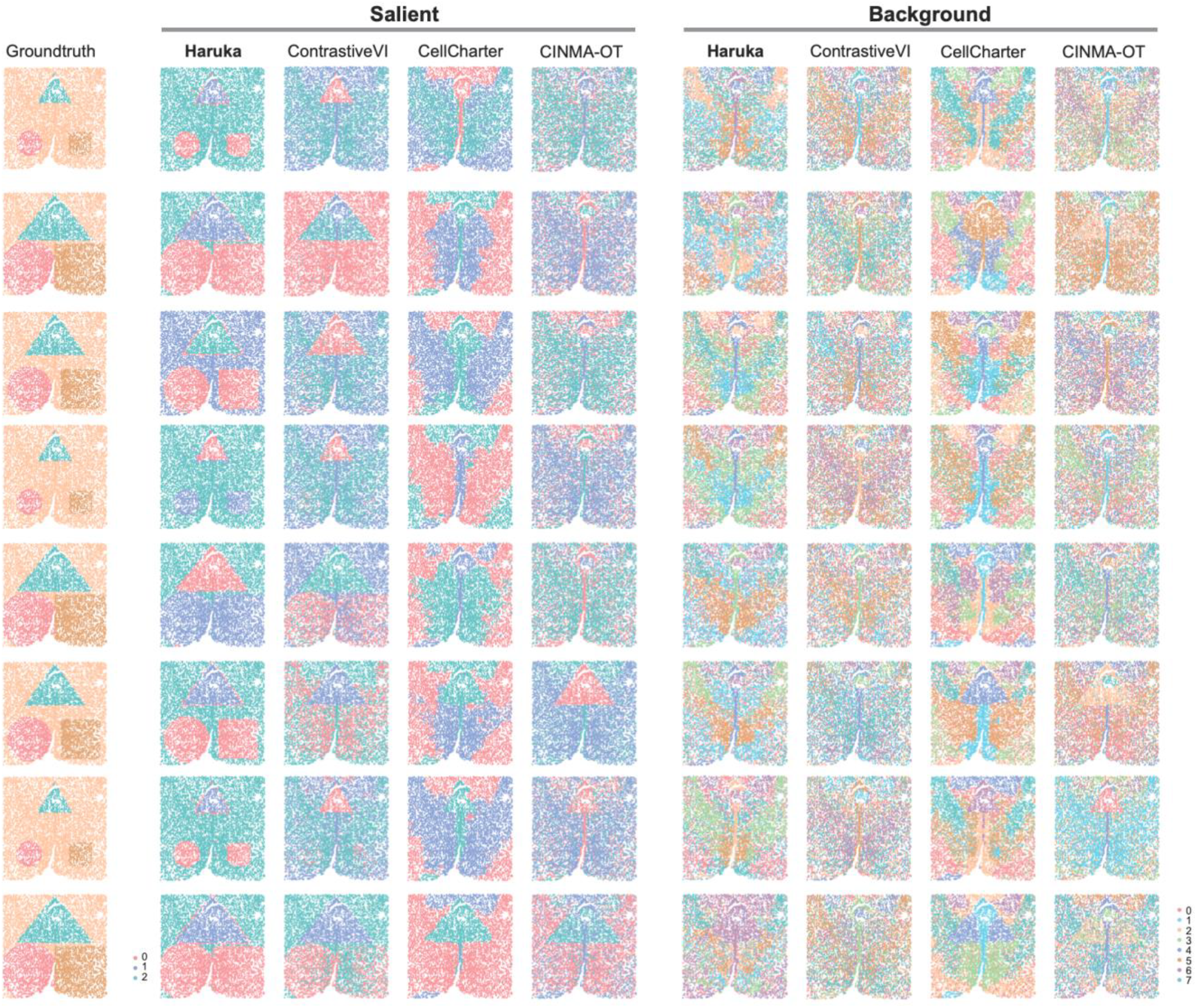
Semi-simulation benchmarking results of Haruka with other comparative methods. Visualization of the ground truth salient and background spatial domains, and the results from Haruka alongside comparison methods (ContrastiveVI, CellCharter, and CINEMA-OT) across different simulated salient shifts (Poisson signal parameter (alpha=2,4,6) and domain area proportion (0.1, 0.15, 0.2)).

**Extended Data Figure 2.**
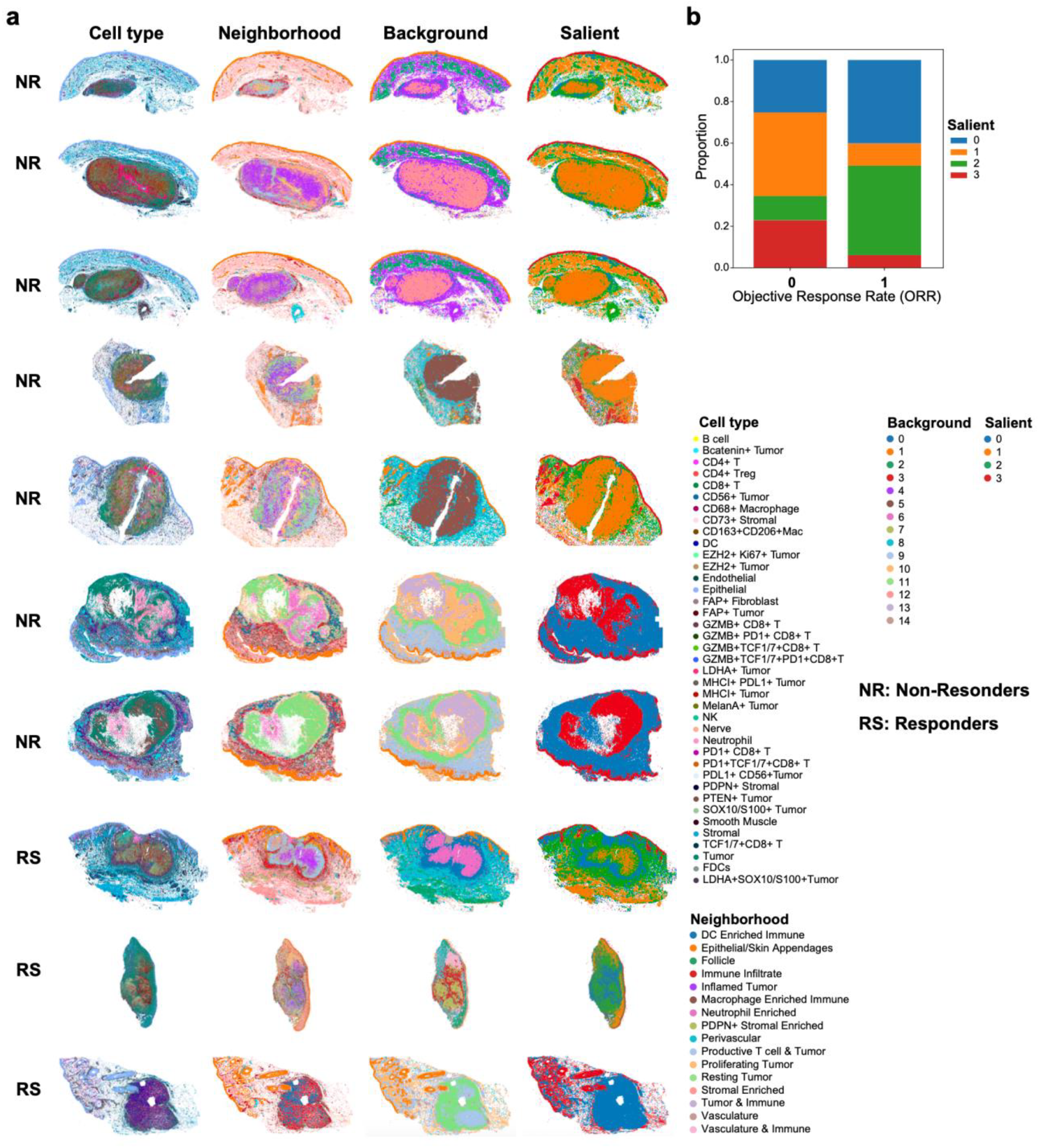
Application of Haruka to the CODEX melanoma immunotherapy dataset. **a**, Spatial visualization of annotated cell types, cell neighborhoods, and Haruka-derived background and salient spatial domains across all samples. **b**, Proportion of salient domains in responder and non-responder groups, showing that salient domains 1 and 3 are predominantly enriched in non-responders.

**Extended Data Figure 3.**
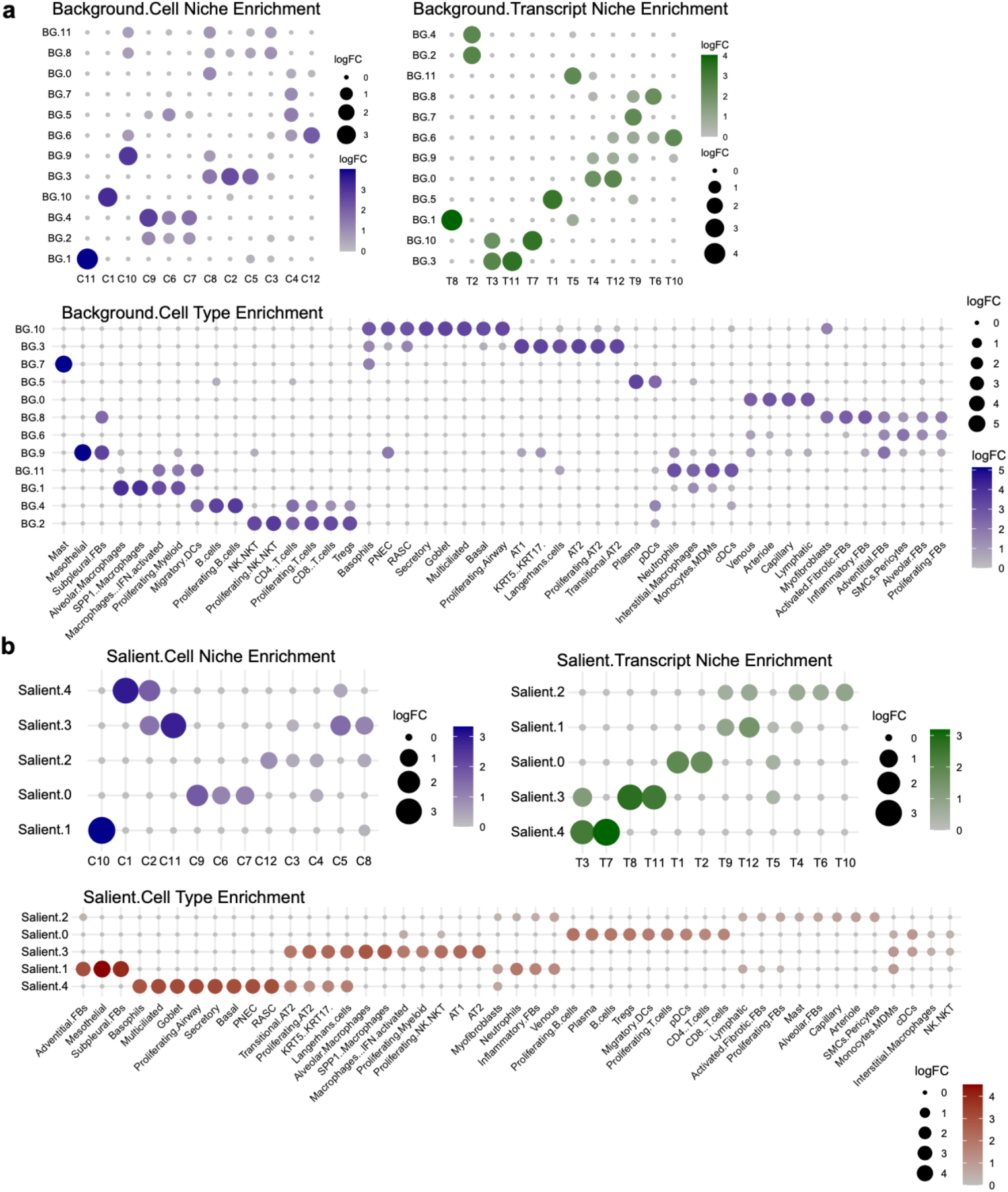
Application of Haruka to the human lung fibrosis 10x Xenium dataset. **a**, Cell niche, transcript niche and cell type enrichment across background spatial domains. Dot size and color denote the log-transformed fold change reflecting enrichment significance. **b**, Cell niche, transcript niche and cell type enrichment across salient spatial domains. Dot size and color denote the log-transformed fold change reflecting enrichment significance.

**Extended Data Figure 4.**
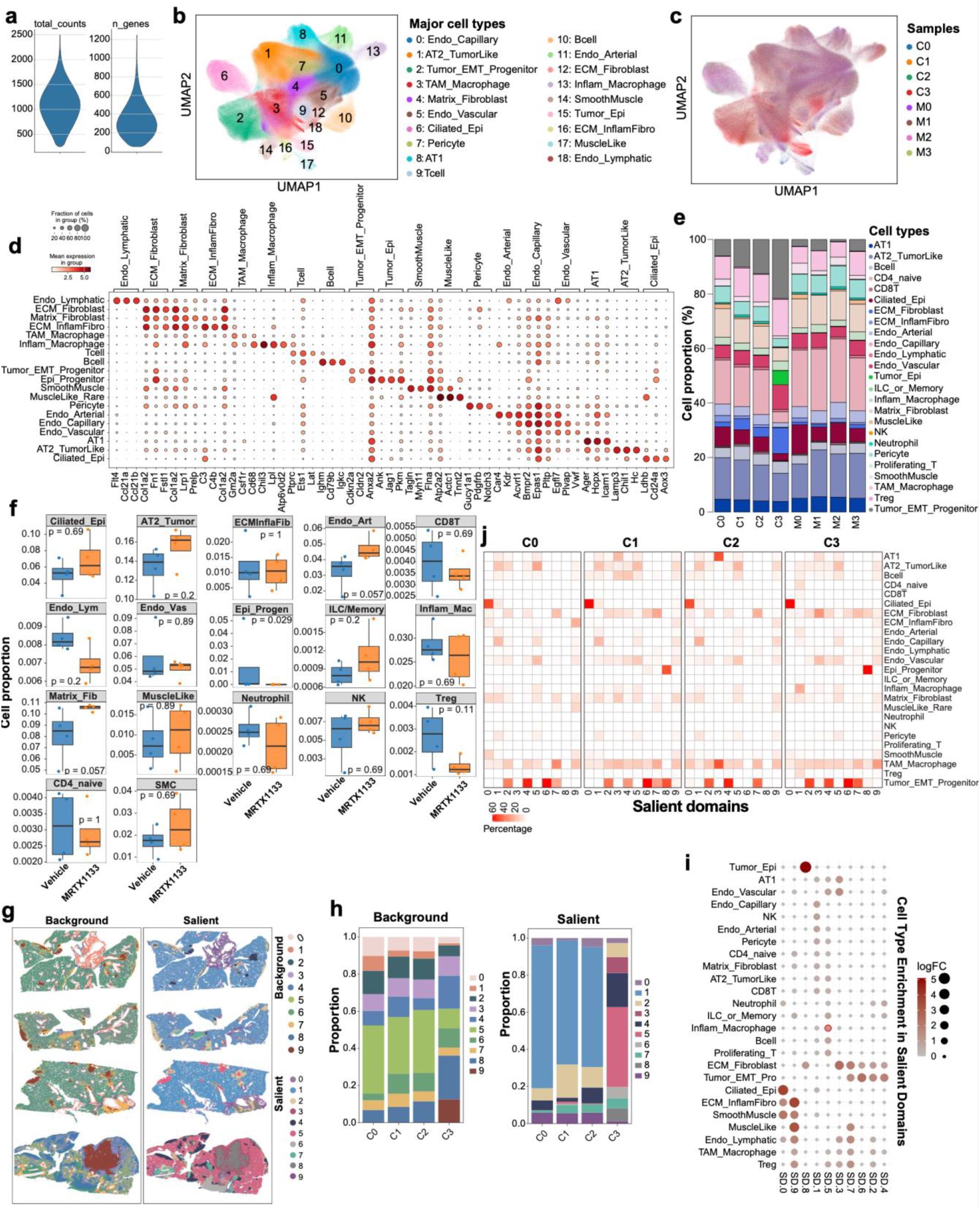
Spatial transcriptomics data processing and application of Haruka in KRAS^G12D^-mutated lung cancer. **a**, Distribution of total transcript counts and number of detected genes per cell in the Xenium dataset. **b-c**, UMAP visualization of major cell type annotations and their distribution across individual samples. **d**, Top three marker genes identified for each major cell type. **e**, Proportions of cell types across all samples. **f**, Comparison of cell type proportions between vehicle- and MRTX1133-treated tumors (n = 4 biological replicates per group). *P*-values were calculated using the Wilcoxon rank-sum test. **g**, Haruka-derived background and salient spatial domains across replicates, revealing distinct microenvironmental organizations within and between treatment groups. **h**, Distribution of background and salient spatial domain proportions across vehicle-treated groups. **i**, Cell type enrichment across salient spatial domains. Dot size and color denote the log-transformed fold change reflecting enrichment significance. Notably, inflammatory macrophages are highly enriched in salient domain 5, highlighted by the red circle. **j**, Cell type proportions across salient domains in vehicle-treated samples, shown per individual tissue slice.

**Extended Data Figure 5.**
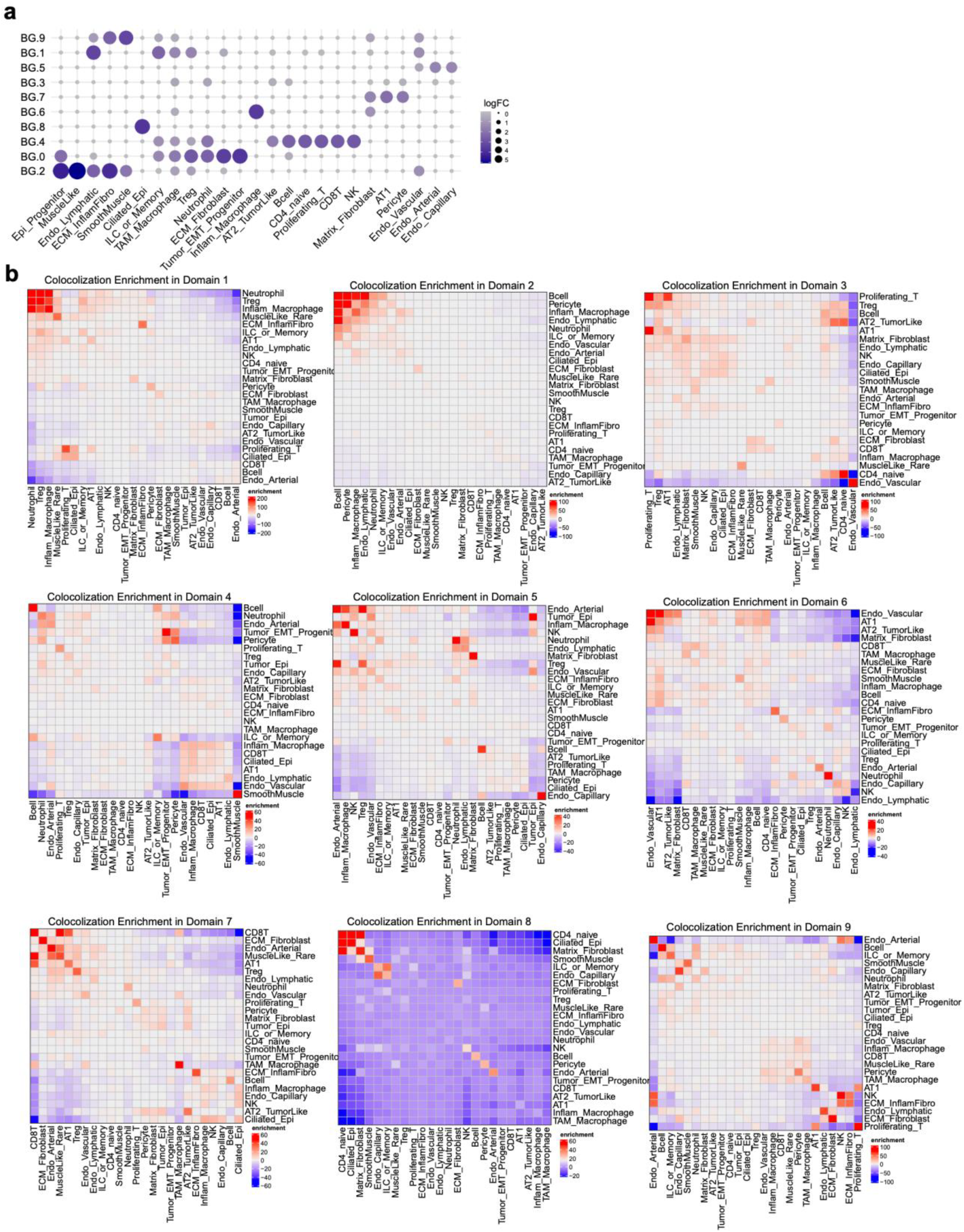
Application of Haruka to investigate the microenvironmental basis of KRAS inhibitor resistance. **a**, Cell type enrichment across background domains, with dot size and color representing log-transformed fold change of enrichment significance. **b**, Domain-specific colocalization enrichment map showing cell-cell proximity patterns in salient domains; higher enrichment values indicate stronger colocalization between corresponding cell-type pairs.

**Extended Data Figure 6.**
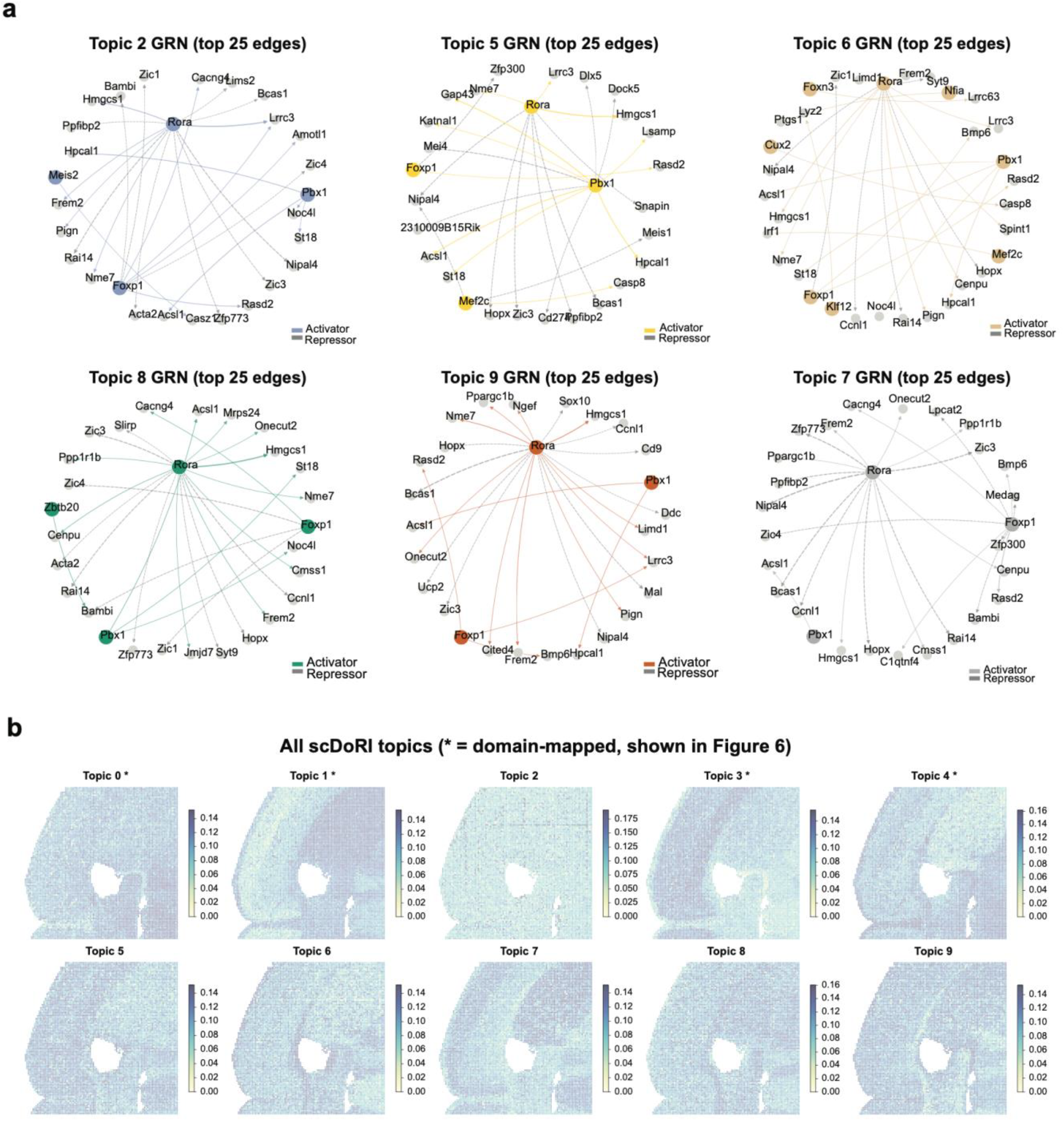
Complete scDoRI topic GRNs and spatial topic maps. **a**, Gene regulatory networks (top 25 edges) for Topics 2, 5, 6, 7, 8, and 9. Recurrent hub TFs include Rora, Foxp1, and Pbx1; shared targets include Bcas1, Hopx, Nipal4, Hmgcs1, and Acsl1. **b**, Spatial maps of all ten scDoRI topic weights. Asterisks denote domain-mapped topics shown in Figure 6 (Topics 0, 1, 3, 4).

**Extended Data Figure 7.**
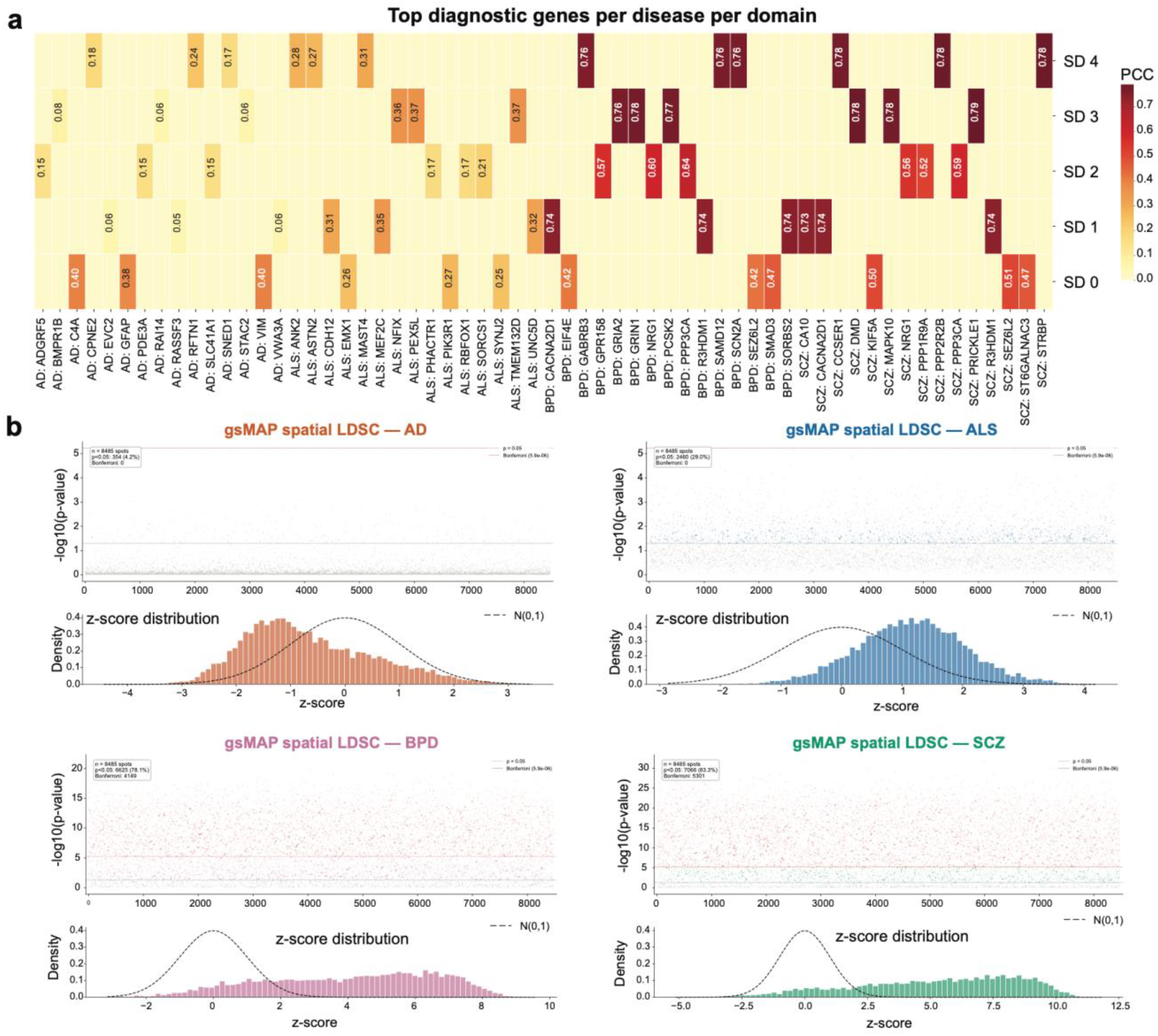
gsMAP diagnostic gene analysis and spot-level GWAS enrichment. **a**, Heatmap of top 15 diagnostic genes per GWAS trait per salient domain, colored by Pearson correlation coefficient (PCC). **b**, Spot-level gsMAP spatial LDSC results for each trait, showing −log_10_(p) per spot with nominal (p = 0.05) and Bonferroni thresholds (top), and empirical z-score distributions against the N(0,1) null (bottom). AD: 4.2% nominal, 0 Bonferroni; ALS: 29.0% nominal, 0 Bonferroni; BPD: 78.1% nominal, 4,149 Bonferroni; SCZ: 83.3% nominal, 5,301 Bonferroni.

## Methods

### Haruka Details

#### Data preprocessing

The input spatial omics dataset was represented as *D* = {**X**,**Y**,**S**}, where **X** ϵ ℝ^*n*⨯*p*^ is the gene expression matrix for *n* cells *p* and genes, 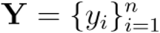 denotes the condition labels for each cell (*y_i_* ϵ { control, condition }), and 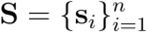 represents the spatial coordinates of cells. Raw gene expression counts were used as input to fit the modeling of the prior gene likelihood distribution.

#### Covariance embedding construction

Haruka adopts the COVET framework ^62^ to encode the spatial niche of each cell. For each cell *i*, the niche is defined as the *k* nearest spatial neighbors based on Euclidean distance in the coordinate space **S**. The gene expression matrix within the niche is denoted E_*i*_ ϵ ℝ^*k*×*g*^, where is *g* the number of selected genes or proteins. To capture the local covariate structure, Haruka computes the shifted covariance matrix Σ_*i*_ as:

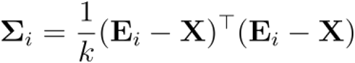

where **X** is the global mean expression vector across the entire dataset, following the shifted covariance approach in scenvi^62^. This global centering ensures that the covariance matrices Σ_*i*_ for different cells are comparable, as they are all constructed relative to the same global mean. The matrix square root of Σ_*i*_, denoted 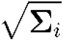, is computed via spectral decomposition. The square root matrix is then vectorized to produce the covariance embedding 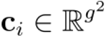. This vector is *c_i_* concatenated with the original expression vector to form the augmented feature vector:

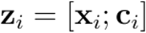

#### Latent variable modeling

The model operates differently for control and condition groups, for condition group cells (*y*_*i*_ = condition), two latent variables are inferred: salient 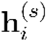 and background 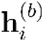

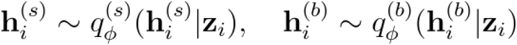

For control group cells (*y*_*i*_ = control), only the background latent variable is inferred:

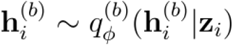

#### Reconstruction loss

Haruka jointly reconstructs gene/protein expression vector x_*i*_ and the covariance embedding vectorc_*i*_ (derived from 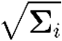). The reconstruction likelihood decomposes as:

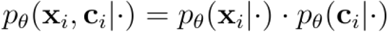

where denotes the corresponding latent variables.

For gene/protein expression/chromatin accessibility (ATAC) data x_*i*_:

- Spatial transcriptomics data: Modeled via a negative binomial distribution:

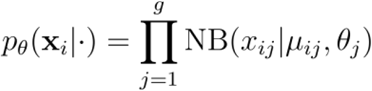
- Spatial proteomics data: Modeled via a Gaussian distribution:

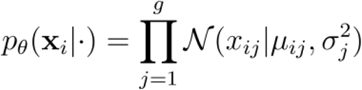
- Spatial chromatin accessibility (ATAC) data: Modeled via a Poisson distribution

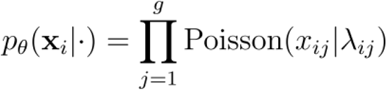

For covariance embeddings c_*i*_:
- Modeled via a Gaussian distribution over the matrix square root:

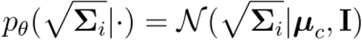

where ***μ***_*c*_ is the predicted mean of the reconstructed matrix square root, and the identity covariance I reflects Euclidean space over matrix square roots. This Gaussian likelihood over 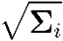 aligns with the AOT metric formulation in scenvi, ensuring computational efficiency. The total reconstruction loss combines log-likelihoods of expression and covariance embeddings:

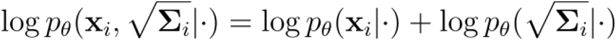

This formulation ensures Haruka reconstructs both the expression-level variations and the covariance structures (niche interactions), in accordance with scenvi’s principled approach.

#### Contrastive alignment and optimization

To ensure disentanglement between salient and background latent spaces, Haruka applies a Wasserstein penalty between background latent distributions of condition (*D*_*t*_) and control (*D*_*b*_) groups:

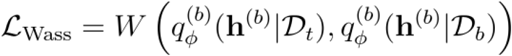

The total loss function is the evidence lower bound (ELBO), which includes the reconstruction of both gene/protein expression and covariance embeddings, combined with the Wasserstein penalty:

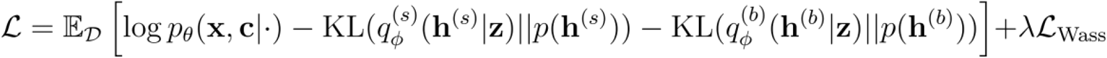

where λ is the regularization coefficient.

#### Spatial neighbor graph construction and latent feature aggregation

Post-training, salient **h**^(*s*)^ and background **h**^(*b*)^latent representations were extracted for target cells. For each tissue slice, a spatial neighbor graph *G*_*S*_ = (*V*_*S*_, *E*_*S*_) was constructed using cell coordinates. Aggregated latent features across *l* = 3 spatial layers were computed as:

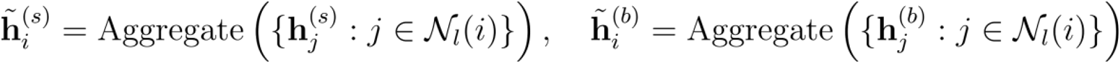

where *N*_l_(*i*) denotes the neighborhood of cell *i* within *l* layers.

#### Identification of spatial domains via Gaussian mixture modeling

Aggregated salient latent representations 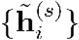 and background latent representations 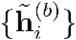 were utilized to define salient spatial domains and background spatial domains, respectively. For each set of aggregated features, a Gaussian Mixture Model (GMM) clustering approach was applied to capture domain structures while modeling cluster-specific covariance:

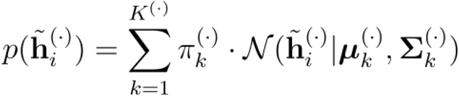

Here, (·) denotes either salient (*s*) or background (*b*) representations. 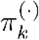 represents the mixing proportion for the *k*-th component, ensuring that 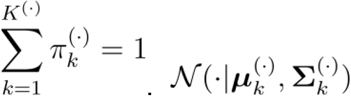. denotes a multivariate normal distribution with mean 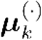 and covariance 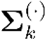. The number of clusters for salient domains *K*^(*s*)^ and background domains *K*^(*b*)^ was predefined (e.g., *K*^(*s*)^ = 4, *K*^(*b*)^ = 8) based on experimental design or model selection criteria. Cluster assignments were determined via maximum a posteriori (MAP) estimation:

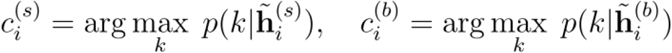

In this context, 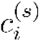 and 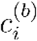 denote the salient and background spatial domain labels for cell *i*, respectively.

### Simulation Dataset and Benchmarking

#### Dataset Overview

To systematically evaluate Haruka’s ability to detect condition-relevant spatial niches under diverse perturbation scenarios, we generated a series of synthetic spatial transcriptomics datasets based on real MERFISH mouse brain data^63^. The original MERFISH slices were used as the base scaffold. Three non-overlapping synthetic spatial domains, corresponding to circle, square, and triangle shapes, were embedded into the tissue space for each dataset. The domain regions covered approximately 5%, 15%, or 25% of the total area, enabling simulation of varying levels of perturbation severity.

Within each spatial domain, non-negative noise was added to the baseline gene expression matrix to simulate perturbation effects. Different Gamma-distributed noise patterns were assigned to each domain, with the circle region perturbed using a Gamma distribution with shape 1 and scale 0.15, the square region using Gamma with shape 1 and scale 0.1, and the triangle region using Gamma noise with shape values of 2, 4, or 6 and a scale of 0.2 across different simulation settings. By varying both the spatial area proportions and noise parameters, multiple simulated datasets were constructed representing distinct perturbation intensities. Each perturbed slice was paired with the corresponding unmodified control slice to form condition-control experimental pairs, with ground truth spatial region annotations recorded for evaluation.

#### Comparison Methods and Metrics

The performance of Haruka was benchmarked against three baseline methods: ContrastiveVI^16^, CInema-OT^17^, and CellCharter^15^. To assess clustering accuracy, we computed the Adjusted Rand Index (ARI) and Normalized Mutual Information (NMI) between the detected salient clusters and the ground truth shape region labels. ARI measures the similarity between predicted clusters and true clusters, adjusting for random chance, and is particularly suitable for evaluating unsupervised clustering methods. Let *n* be the total number of cells, *n*_*ij*_ the number of cells in both predicted cluster and true cluster *j, a*_*i*_ the sum over row *i*, and *b*_*j*_ the sum over column *j* in the contingency table:

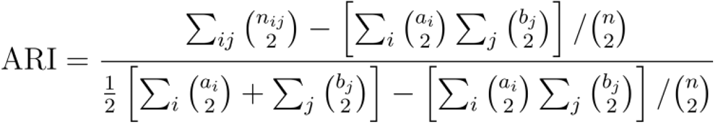

NMI quantifies the mutual dependence between cluster assignments and ground truth, providing a normalized score that accounts for differences in cluster size. Let *C* and *G* be the sets of predicted and ground truth clusters, respectively, and let *I*(*C*; *G*) denote mutual information and *H* (*C*), *H*(*G*)denote their entropies:

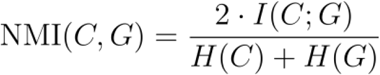

where:

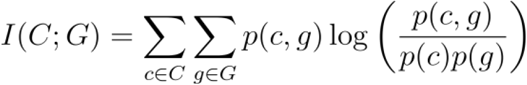

and

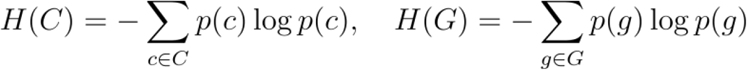

In addition to clustering accuracy, we evaluated the spatial coherence of the detected domains by computing CHAOS and PAS scores. CHAOS measures local spatial smoothness based on nearest-neighbor label consistency For each cluster *k*, we compute the average pairwise Euclidean distance between all cells *x*_*i*_, *x*_*j*_, ∈ *C*_*k*_ within the same cluster:

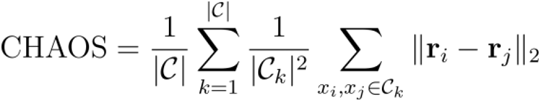

where r_*i*_ denotes the spatial coordinates of cell *x*_*i*_ and |*C*_*k*_| is the size of cluster *k*.

While PAS captures global spatial autocorrelation of cluster labels.

Let *x*_*i*_ be a cell with spatial neighbors *N*(*x*_*i*_), and *ŷ*_*i*_ its cluster label. PAS is defined as:

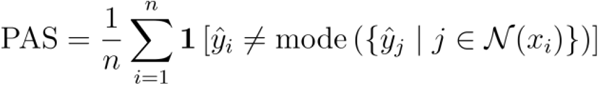

Where **1**[·] is the indicator function and mode denotes the most frequent label among neighbors.

Lower CHAOS and PAS scores indicate better spatial continuity and less fragmentation. Together, these four metrics comprehensively evaluate both the correctness of salient cluster assignments and the preservation of underlying spatial structure.

### Benchmarking on Mouse Brain Aging Dataset

#### Dataset Overview

To further evaluate the performance of Haruka and baseline methods in real-world scenarios, we conducted benchmarking experiments on a large-scale spatial transcriptomics dataset profiling mouse brain aging^8^. Specifically, we utilized the spatial transcriptomic data generated in spatial aging clock dataset, which profiled the mouse brain across 20 different age points throughout the lifespan using MERFISH technology. For our benchmarking, we selected slices corresponding to embryonic day 3.4 (E3.4) and embryonic day 34.5 (E34.5) as the control and condition groups, respectively. This selection reflects early and mid-gestational brain developmental stages, allowing us to assess spatially driven aging responses under well-separated biological conditions.

#### Label generation of response heterogeneity

We generated two types of ground-truth labels. Background domain labels were defined based on anatomical subregions annotated in the original study using an unbiased approach. These subregions, considered relatively unaffected by aging, were designated as background domains. Salient domain labels were derived from previously reported correlations between cell types and aging across different brain sections (CTX, STR, VEN, and CC/ACO), based on the full 20-timepoint aging timeline provided in the original study (Extended Data). For each brain section, we identified cell types that were significantly positively correlated with age (Pearson correlation > 0.4, p < 0.05), negatively correlated with age (Pearson correlation < −0.4, p < 0.05), or uncorrelated. This allowed us to define three salient categories per brain section—positive, negative, and uncorrelated—resulting in a total of 12 salient domains (3 per section × 4 sections). For background domains, we used the fine-grained anatomical annotations from the original study, which were generated in an unbiased manner and remained stable throughout the aging process. These subregions served as reliable indicators of the spatial background.

#### Comparison Methods and Metrics

We adopted the same baseline methods used in the simulation benchmark—CellCharter, ContrastiveVI, and CINEMA-OT—with identical implementation pipelines, clustering algorithms (GMM), and target cluster numbers to ensure fair comparisons. For evaluation, we reported ARI and NMI metrics only, excluding spatial continuity metrics (CHAOS, PAS). This exclusion was motivated by the nature of our ground-truth domains, which include both spatially contiguous regions and small, spatially disconnected niches. These small niches are particularly important due to their difficulty of detection within the same anatomical subregions, and continuity-based metrics would undervalue their significance.

By using this biologically motivated, region-based ground truth and spatially relevant evaluation framework, we aimed to rigorously assess the ability of Haruka and comparative methods to capture salient spatial responses to aging in complex, heterogeneous brain tissue.

### Haruka application on Melanoma Immunotherapy CODEX Dataset

#### Dataset Overview

We applied Haruka to a human metastatic melanoma dataset comprising 12 FFPE tumor samples from six patients with Stage IV disease^8^. Patients received checkpoint inhibitor therapy, with samples harvested at pre-treatment (baseline) and post-treatment timepoints. Tumor tissues were profiled via CODEX multiplexed imaging (58-antibody panel), yielding 5,019,159 segmented cells across immune, stromal, and tumor compartments. In our experiment, we utilized unsupervised clustering to identify 39 major cell types, setting post-treatment as the condition sample and pre-treatment as the control group. In our experiment, we set post-treatment as the condition sample and pre-treatment as the control group

#### Integration of Salient Spatial Domains with scRNA-seq Using MaxFuse

To explore downstream utility of Haruka’s salient spatial domains, we performed a cross-modality integration between protein-based spatial domains derived from CODEX imaging and unpaired single-cell RNA sequencing (scRNA-seq) profiles. Specifically, we aimed to assess whether Haruka-identified domains could act as spatial anchors to enrich scRNA-seq data with spatial context and uncover gene expression programs associated with immunotherapy outcomes in melanoma.

We first identified salient domains in CODEX data using Haruka. Among these, salient domain 4 was significantly enriched in samples with no objective response rate to immunotherapy. We thus designated this as the immunotherapy-resistant reference domain. Cells assigned to this domain were used as the “reference anchor” for downstream MaxFuse integration.

To link unpaired scRNA-seq data with this CODEX-defined spatial anchor, we employed MaxFuse^64^, a modality-agnostic matching algorithm that iteratively aligns two datasets using both cell-cell similarity and cross-modality feature correlation. The scRNA-seq data consisted of a post-immunotherapy melanoma dataset profiling T cell population. We performed feature preprocessing separately for each modality: the CODEX protein panel was normalized via arcsinh transformation, while scRNA-seq gene expression matrices were log-normalized and filtered to retain highly variable genes overlapping with markers in the CODEX panel. The integration was performed with MaxFuse’s default pipeline, which iteratively refines cross-modal similarity through canonical correlation analysis (CCA)-based latent space alignment, followed by nearest neighbor bipartite matching and denoising propagation.

From this process, we obtained one-to-one correspondences between scRNA-seq cells and Haruka/CODEX-defined spatial regions. In particular, scRNA-seq cells matched to salient domain 4 were labeled as “poor immunotherapy responders,” and the rest were considered the control group. We restricted the analysis to CD8^+^ T cells, based on scRNA-seq cell type annotation, and investigated transcriptional differences between these two subgroups.

#### Differential Expression and Gene Set Enrichment Analysis

To characterize molecular signatures associated with immunotherapy resistance, we performed differential gene expression (DE) analysis between the CD8^+^ T cells assigned to the non-responder salient domain and the remaining CD8^+^ T cells. The DE was performed using a Wilcoxon rank-sum test implemented in Scanpy, with Benjamini– Hochberg correction applied to control false discovery rate. Genes with adjusted *p*-value < 0.05 and |log2 fold change| > 0.25 were retained for downstream enrichment. We then conducted Gene Set Enrichment Analysis (GSEA) on the ranked DE gene list using the *gseapy* implementation with the GO Biological Process database. We used adjusted p-value < 0.05 as cutoff for significant ones.

#### Multimodal Association Analysis

To quantify the cross-modality association between a protein marker *p* and a matched gene expression profile *g*, we computed their Pearson correlation coefficient across aligned cells in a given spatial domain. Let *p* = {*p*_1_, … *p*_*n*_} denote the protein intensity vector across CODEX cells, and *g* = {*g*_1_, … *g*_*n*_} the RNA expression vector across the matched scRNA-seq cells.

The correlation is defined as:

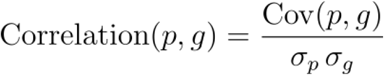

where is Cov (*p,g*) the covariance between protein and gene signals, and *σ_p_, σ_g_* are the standard deviations of *p* and *g*, respectively.

This metric reflects the strength of linear association between CODEX protein signals and RNA expression patterns within aligned spatial domains, with values ranging from −1 (perfect negative correlation) to (perfect positive correlation).

### Haruka application on Lung Fibrosis Xenium Dataset

#### Dataset Overview

We analyzed spatial transcriptomics data from 45 human lung samples^45^ (26 pulmonary fibrosis, 9 healthy controls) using the Xenium platform with a 343-gene panel. The dataset included 1.6 million cells, annotated into 12 spatial niches (C1-C12) reflecting fibrotic, inflammatory, and epithelial regions.

#### Alignment Benchmarking with SLAT Using Haruka-Derived Representations

To evaluate the fidelity of spatial alignment across fibrotic progression, we applied SLAT, a spatial domain alignment method, to paired tissue slices representing less affected and more affected fibrotic states from a human lung fibrosis spatial transcriptomics dataset. These samples were drawn from a publicly available Xenium-based spatial profiling study, where histopathological assessment had previously stratified regions according to the severity of fibrosis. The evaluation focused on whether Haruka’s learned background domain representations—designed to isolate stable, condition-invariant features—could enhance alignment across varying tissue conditions.

We used two types of input representations for SLAT^43^: the original gene expression matrices and Haruka-derived background domain embeddings. The background embeddings were obtained through Haruka’s contrastive variational inference framework, which is trained to separate shared, condition-invariant structure from perturbation-specific effects by learning latent representations that reflect stable spatial context. Specifically, Haruka learns background latents solely from control (less affected) conditions, thus retaining architectural consistency while removing condition-induced gene expression variation.

SLAT was then run independently on each pair of slices, once with the original gene expression data and once with the Haruka background representations. Its alignment procedure incorporates both spatial proximity and transcriptomic similarity using a graph-based latent optimization framework, resulting in slice-to-slice alignment of spatial domains. To benchmark alignment quality, we used CNiche domain annotations from the original study as unbiased ground truth references for architectural domains. The performance was quantified using two metrics, F1-score and accuracy.

Formally, Let *S*(*a*) and *S*(*b*) denote two spatial transcriptomics slices (e.g., less affected and more affected tissue sections), containing *n*_*a*_ and *n*_*b*_ cells, respectively. Each cell is represented by a feature vector—either the original gene expression or a lower-dimensional embedding (e.g., Haruka background representation).

The SLAT model takes the two slices and outputs a matching matrix

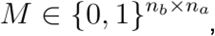

where

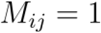

indicates that cell *I* in slice *S*(*b*) is matched to cell *j* in slice *S*(*a*).

Each row of *M* contains exactly one non-zero entry, corresponding to the one-to-one matched reference cell.

Let 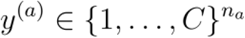 be the ground truth CNiche domain labels for slice *S*(*a*).

Using the matching matrix *M*, we define the predicted label for each cell *S*(*b*) in by propagating the label of its matched counterpart:

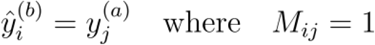

This allows us to assign predicted CNiche labels 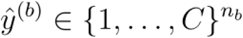 to all cells in the target slice *S*(*b*).

To evaluate alignment quality, we compare the predicted labels *ŷ*^(*b*)^to the ground truth labels *y*^(*b*)^ from the original annotation of *S*(*b*).

We here utilize three metrics to evaluate the match situation between predicted labels *ŷ*^(*b*)^to the ground truth labels *y*^(*b*)^. The micro-averaged F1-score is computed globally by counting the total true positives (TP), false positives (FP), and false negatives (FN) across all CNiches by the matched cell CNchihe as prediction and ground truth CNiche as labels:

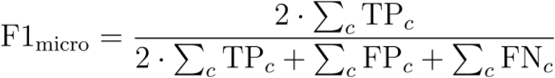

The macro-averaged F1-score is calculated by first computing the F1-score for each class independently and then taking their unweighted average:

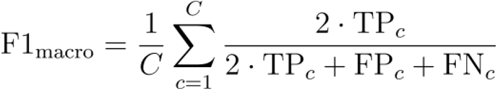

Where *C* is the number of CNiche classes.

Accuracy quantifies the overall proportion of correctly matched cells among all cells:

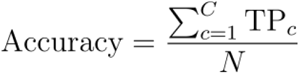

Where *N* is the total number of cells s and TP_*c*_ refers to the number of correctly matched cells for CNiche *c*.

#### Cell State Transition Stratification

Beyond evaluating alignment performance, we sought to leverage Haruka-SLAT alignment to dissect early molecular events that occur during fibroblast transition in fibrosis. Using aligned slices in the Haruka background space, we identified matched pairs of alveolar fibroblasts (Alv-FBs) between less affected and more affected conditions. In the less affected slice, Alv-FBs were then stratified based on whether their SLAT-aligned counterparts in the more affected slice retained an alveolar identity or had transitioned into an activated fibrotic state. This matching defined two subpopulations within the phenotypically similar Alv-FBs in the less affected tissue: one maintaining a stable identity and one primed for activation. Despite similar spatial localization and cell-type identity, the primed subgroup exhibited gene expression features suggestive of early transition toward fibrotic activation. This observation was leveraged to explore pre-fibrotic transcriptional signatures that might serve as early indicators of disease trajectory.

#### Differential Expression and Gene Ontology Analysis

To identify early molecular changes predictive of fibroblast activation, we performed differential gene expression analysis between the stable and primed Alv-FB subgroups using the Wilcoxon rank-sum test. Genes with a Benjamini–Hochberg corrected *p*-value below 0.05 and an absolute log2 fold change greater than 0.25 were considered significant. The resulting gene list was subjected to Gene Ontology enrichment analysis using the Enrichr^65,66^ module from the gseapy package. Enriched biological processes in the primed population included extracellular matrix organization, cytoskeletal remodeling, and cell adhesion—processes closely linked to early fibroblast activation and fibrogenesis. These results suggest that even in structurally preserved regions, Haruka’s background-based matching can reveal molecular precursors of pathogenic state transitions.

### Haruka analysis of KRAS-inhibitor Xenium 5K dataset

#### Mouse Model and Ethics Statement

All animal experiments were performed in accordance with institutional guidelines and approved by the Committee on Animal Care (IACUC for MIT and Whitehead Institute). Mice were housed in the Whitehead Institute Animal Facility under a 12-hour light/dark cycle with ad libitum access to food and water. C57BL/6J Albino (Jackson Labs strain # 000058) female mice aged 8 weeks were used for experimentation

#### Tumor Cell Preparation and Engraftment

A murine KRAS^G12D^-mutant lung adenocarcinoma cell line (KP1233, provided as a gift from Dr. Tyler Jacks” lab, Koch Institute for Integrative Cancer Research at MIT) was cultured in 1X DMEM (ThermoFisher Gibco™ Cat#13345364) supplemented with 10% heat-inactivated (56°C for 30min) ultra-low IgG FBS (ThermoFisher Gibco™ Cat# 16250078) under standard conditions (37°C, 5% CO_2_). Cells were tested for individual murine pathogens (IMPACT I, Corynebacterium bovis, Mycoplasma pulmonis). Cells were harvested at ~70–80% confluency using TrypLE Express (ThermoFisher), washed with complete media, and resuspended in PBS at a final concentration of 1 × 10^6 cells per 100 µL. For establishment of lung metastases, mice were injected via the lateral tail vein with 100 µL of the tumor cell suspension. Mice were monitored daily for signs of distress and body weight changes.

#### Treatment

Seven days post-injection, after confirmation of tumor establishment (based on prior model optimization), mice were randomized into two treatment groups: vehicle control (formulation matching MRTX1133 vehicle); MRTX1133 (KRAS G12D inhibitor; MedChemExpress Cat# HY-134813; dissolved in 10% Captisol and 50mM citrate buffer pH 5.0). MRTX1133 or vehicle was administered by intraperitoneal injection once daily for 5 consecutive days at a dose of 30 mg/kg, as described in Hallin et. al^67^.

#### Tissue Collection, Processing and Xenium Experiment

At the experimental endpoint (day 21 post-injection), mice were euthanized by CO_2_ inhalation followed by cervical dislocation. Lungs were dissected, visually inspected for tumor burden, and processed. The lung tissues were inflated with fixative (10% formalin) and processed according to the manufacturer’s protocol for the Xenium In Situ platform (10x Genomics). Samples were embedded, sectioned, and mounted such that each slide contained 4 distinct tumor-bearing lung samples from individual mice (n = 4 biological replicates per condition). For spatial profiling, each condition was assigned to a single Xenium slide containing multiple lung samples from different mice. All sample processing and library preparation were performed in parallel to minimize batch effects.

#### Data Generation Procedure and Preprocessing

Raw Xenium outputs were processed using the Xenium Onboard Analysis (XOA) pipeline (10x Genomics), which performs image alignment, spot detection, molecule calling, and cell segmentation based on integrated nuclear and membrane stains. Segmentation-derived cell boundaries were quality-checked to remove artifacts, doublets, and low-confidence cells. Gene count matrices were then aggregated across individual lung samples and converted into an AnnData object for downstream analyses. To harmonize data across slides and biological replicates, we applied per-cell quality control filters, excluding cells with extremely low molecule counts (< 50) or high mitochondrial read fractions (> 5%)^68,69^. Gene expression matrices were normalized to account for sequencing depth, followed by log-transformation. For visualization and exploratory analyses, we generated low-dimensional embeddings using principal component analysis (PCA) and Uniform Manifold Approximation and Projection (UMAP).

#### Cell Type Annotation

Following dimensionality reduction, we performed unsupervised clustering using the Leiden algorithm on the neighborhood graph computed from the top 40 principal components (Scanpy’s sc.pp.neighbors, sc.tl.leiden, resolution = 0.6 unless otherwise specified). The resulting clusters were visualized using UMAP and inspected for biological coherence based on known cell type markers. To annotate clusters, we identified differentially expressed genes (DEGs) using the rank_genes_groups function in Scanpy with the Wilcoxon rank-sum test. For each cluster, we extracted the top 50 marker genes and compared them to canonical cell type markers from both lung and immune cell atlases as well as prior publications on KRAS-driven lung cancer models. To resolve T cell heterogeneity, we applied additional Leiden clustering within the T cell compartment to identify finer subtypes. Final cell type annotations were determined by integrating clustering results, marker gene enrichment, and prior biological knowledge. Populations with ambiguous or transitional features (e.g., tumor-associated fibroblasts, proliferative immune cells) were annotated using descriptive qualifiers to reflect their functional or spatial context.

#### Pathway Activity Score Calculation

We calculated the pathway activity score for cells annotated as tumor cells (“Tumor EMT Progenitor,” “AT2 TumorLike”) using the drug2cell package. The drug2cell^65^ package accepts whole gene expression as input and calculates the activity score for a given gene set in each individual cell (a higher score indicates higher gene set activity in that cell). We utilized the KEGG “Non-small cell lung cancer” and “MAPK signaling pathway” as input gene sets for the drug2cell score calculation. We performed this analysis only on tumor cells to eliminate noise from other cell types.

#### Cell-Cell Communication Analysis

We utilized the LIANA^70^ package with the CellPhoneDB^71^ database for the cell types Treg, and Proliferating T cells within salient domain 0. LIANA infers cell-cell communication by calculating the mean ligand and receptor gene expression values for each group based on the input data. We used the full panel gene expression as input and selected the top 30 ligand-receptor (LR) pairs for downstream analysis. We used CCPlotR^72^ for visualization.

### Haruka analysis of spatial multi-omics in mouse brain

#### Spatial ATAC–RNA-seq Dataset and Preprocessing

We obtained paired spatial ATAC–RNA-seq data from the spatiotemporal tri-omic atlas of the postnatal mouse brain ^51^, comprising two P21 C57BL/6 mouse brain sections: an uninjured control (P21S3) and a section subjected to focal demyelination by stereotactic lysolecithin (LPC) injection into the corpus callosum (LPC21S1). RNA count matrices were filtered to in-tissue spots with standard QC (≥10 cells per gene, ≥10 genes per cell) using Scanpy. ATAC fragments were processed via SnapATAC2 against the mm10 genome, and a peak-by-cell matrix was built against ENCODE reference peaks with paired-insertion counting, removing peaks detected in fewer than 1% of cells. Only barcodes present in both modalities were retained for downstream analysis.

#### Haruka Multimodal Spatial Domain Identification

Haruka was applied to identify condition-specific (salient) and shared (background) spatial domains from paired spatial RNA and ATAC modalities. The top 4,000 highly variable genes and 15,000 highly variable peaks were selected (Seurat v3 flavor), and spatial covariance structure was captured via COVET for each modality. Cell latent embedding encoders were trained independently per modality with the control sample as contrastive background and the LPC sample as target, yielding modality-specific salient and background latent representations 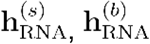, and 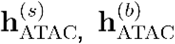. The RNA and ATAC latents were then concatenated to form joint multimodal representations:

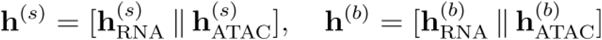

which were subsequently aggregated across *l* = 3 spatial neighborhood layers following the same spatial neighbor graph construction scheme described above, producing niche-level salient and background embeddings 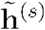 and 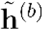. These were clustered to define condition-specific (salient) and shared (background) multimodal spatial domains.

#### scDoRI Topic Modeling and GRN Inference

To infer enhancer-mediated regulatory programs underlying each salient domain, we applied scDoRI^52^ to the paired RNA and ATAC profiles using 4,000 genes, 300 TFs, and 50,000 peaks, with CisBP motif annotations and a ±80 kb peak–gene window. The model learned 10 regulatory topics in a two-phase training scheme (topic model followed by GRN inference), and GRN significance was assessed by 1,000 permutations (p < 0.05). Domain-specific topics were identified by computing z-scored mean topic activation across cells within each salient domain.

#### Genetically Informed Spatial Trait Mapping with gsMAP

To link the multimodal spatial domains to human complex trait genetics, we applied gsMAP^57^ in quick mode, using Haruka’s salient multiomics embedding as the pre-computed latent representation together with the full RNA count matrix, salient domain annotations, and a mouse-to-human homolog mapping. Spatial LDSC was performed against the 1000 Genomes EUR Phase 3 LD reference for four GWAS traits: Alzheimer’s disease, amyotrophic lateral sclerosis (ALS), bipolar disorder (BPD), and schizophrenia (SCZ). Domain-level significance was aggregated via the Cauchy combination test, and top diagnostic genes driving domain-specific enrichment were extracted from the latent-to-gene output.

## Computational Resource

All experiments were performed on a server running Ubuntu 20.04 with a 72-core Intel(R) Xeon(R) CPU E5-2697 v4 @ 2.30GHz and an Nvidia A100 (80 G) GPU.

## Data Availability

MERFISH mouse brain aging data can be downloaded from https://zenodo.org/records/13883177. The CODEX melanoma immunotherapy dataset is available at https://datadryad.org/dataset/doi:10.5061/dryad.k0p2ngfcc. The Xenium lung fibrosis dataset can be accessed through GEO (GSE250346). The KRAS^+^ Xenium cancer dataset is available upon request from the corresponding author. The spatial multi-omics dataset is available at https://www.ncbi.nlm.nih.gov/geo/query/acc.cgi?acc=GSE308623

## Code Availability

All code required to reproduce the figures, results, and tutorials is publicly available at https://github.com/C0nc/Haruka.

## Acknowledgements

We thank the Whitehead Institute Animal Facility for expert assistance with mouse husbandry and for maintaining high-quality experimental conditions. We are grateful to the Broad Institute’s Spatial Technology Platform for support with spatial transcriptomics experiments. This work was supported by the Whitehead Institute and the Whitehead Innovation Initiative. N.S. is supported by the Whitehead Institute Fellows Program, the Whitehead Innovation Initiative, and the NIH Director’s Early Independence Award (DP5OD042638). K.W. is supported by the Valhalla Foundation, AACR-AstraZeneca Career Development Award for Physician-Scientists in Honor of José Baselga, an anonymous grant, Richard Reisman in honor of Jane Reisman and Lilian Reisman, Department of Defense (W81XWH2210141), National Cancer Institute (R01CA279259), the Research Foundation for the Treatment of Ovarian Cancer, ALK Positive Lung Cancer Foundation, and Torrey Coast Foundation. The content of this manuscript is solely the responsibility of the authors and does not necessarily represent the official views of the NIH or other funding agencies.

## Author contributions

Y.C. and N.S. conceived and designed the study. N.S. supervised the overall project. J.B. and K.W. established and implemented the mouse lung cancer model experiments; J.B. performed in vivo studies and sample processing, with experimental supervision from K.W.. Y.C. conducted data analysis and developed the computational modeling framework. K.W. provided scientific input throughout the study.

J.B. and K.W. wrote the experimental methods pertaining to lung cancer. Y.C. and N.S. wrote and revised the manuscript with input from all authors.

